# DOC2b enrichment mitigates proinflammatory cytokine-induced CXCL10 expression by attenuating IKKβ and STAT-1 signaling in human islets

**DOI:** 10.1101/2024.12.22.629540

**Authors:** Diti Chatterjee Bhowmick, Miwon Ahn, Supriyo Bhattacharya, Arianne Aslamy, Debbie C. Thurmond

**Author notes:** Corresponding authors Debbie C. Thurmond, Ph.D., Department of Molecular and Cellular Endocrinology Arthur Riggs Diabetes and Metabolism Research Institute City of Hope Beckman Research Institute, 1500 E. Duarte Road Duarte, CA 91010 USA, Diti Chatterjee Bhowmick, Ph.D., Department of Molecular and Cellular Endocrinology Arthur Riggs Diabetes and Metabolism Research Institute City of Hope Beckman Research Institute, 1500 E. Duarte Road Duarte, CA 91010 USA.

## Abstract

**Introduction:** Type 1 diabetic human islet β-cells are deficient in double C 2 like domain beta (DOC2b) protein. Further, DOC2b protects against cytokine-induced pancreatic islet β-cell stress and apoptosis. However, the mechanisms underpinning the protective effects of DOC2b remain unknown.

**Methods:** Biochemical studies, qPCR, proteomics, and immuno-confocal microscopy were conducted to determine the underlying protective mechanisms of DOC2b in β-cells. DOC2b- enriched or-depleted primary islets (human and mouse) and β-cell lines challenged with or without proinflammatory cytokines, global DOC2b heterozygous knockout mice subjected to multiple-low-dose-streptozotocin (MLD-STZ), were used for these studies.

**Results:** A significant elevation of stress-induced CXCL10 mRNA was observed in DOC2b- depleted β-cells and primary mouse islets. Further, DOC2b enrichment markedly attenuated cytokine-induced CXCL10 levels in primary non-diabetic human islets and β-cells. DOC2b enrichment also reduced total-NF-κB p65 protein levels in human islets challenged with T1D mimicking proinflammatory cytokines. IKKβ, NF-κB p65, and STAT-1 are capable of associating with DOC2b in cytokine-challenged β-cells. DOC2b enrichment in cytokine-stressed human islets and β-cells corresponded with a significant reduction in activated and total IKKβ protein levels. Total IκBβ protein was increased in DOC2b-enriched human islets subjected to acute cytokine challenge. Cytokine-induced activated and total STAT-1 protein and mRNA levels were markedly reduced in DOC2b-enriched human islets. Intriguingly, DOC2b also prevents ER-stress-IKKβ and STAT-1 crosstalk in the rat INS1-832/13 β-cell line.

**Conclusion:** The mechanisms underpinning the protective effects of DOC2b involve attenuation of IKKβ-NF-κB p65 and STAT-1 signaling, and reduced CXCL10 expression.

**Graphical abstract:** 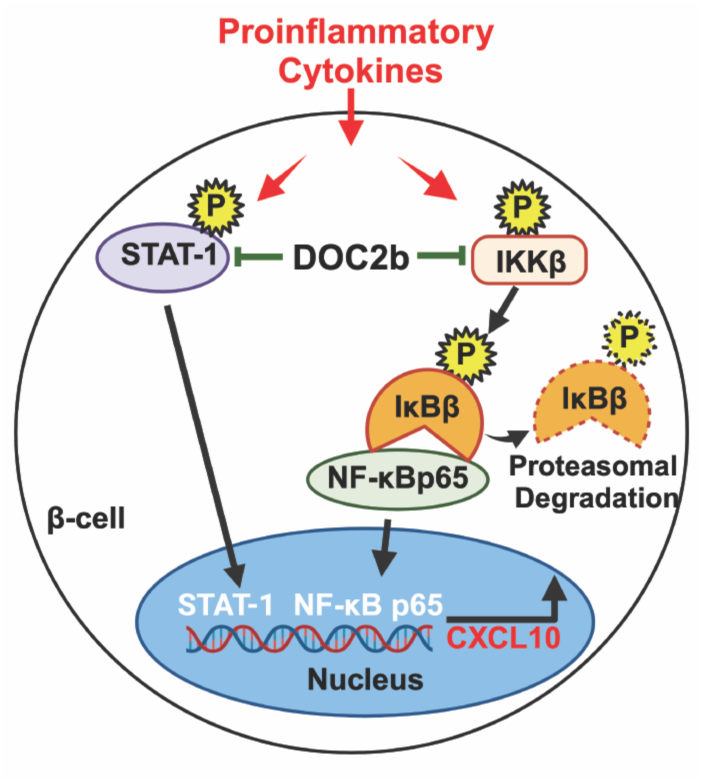

## 1. INTRODUCTION

Type 1 diabetes (T1D) is characterized by dysfunction and autoimmune destruction of the insulin-producing β-cells of the pancreatic islets, resulting in a requirement for life-long insulin replacement therapy. Although the exact cause of T1D is yet to be known, genetic and environmental factors are suggested to play important causal roles. Loss of functional β-cell mass can progress for years before the appearance of detectable T1D symptoms and resulting growing hyperglycemia damages major organs like the heart, blood vessels, nerves, eyes, and kidneys. Mounting evidence indicates that β-cells rather than just being passive targets, are active participants along with the immune system in T1D pathogenesis [1–4]. Thus, curative strategies must provide support for β-cell function as well as for protection from autoimmune destruction.

During T1D, proinflammatory cytokines (e.g., IL-1β, TNFa, and IFNg) released from activated macrophages and T cells induce β-cell destruction. The transcription factor nuclear factor kappa-B (NF-κB) plays a crucial role in mediating IL-1β- and TNFα-induced β-cell dysfunction and apoptosis [5, 6], whereas IFNγ signaling is thought to be activated by STAT-1 [7, 8]. In β-cells, IL-1b and IFN-γ synergistically induce the gene expression and promoter activity of the chemokine ligand CXCL10 via NF-κB p65 and STAT-1 signaling [9]. CXCL10 is produced by immune cells and β-cells, and recruits auto-aggressive lymphocytes to the pancreatic islets during T1D [10–12]. Elevated CXCL10 serum levels are detected in T1D patients [11, 13] and in individuals at high risk for developing T1D [11, 13]. Further, nonobese diabetic (NOD) mice, a surrogate model for human T1D [14], produce CXCL10 in pancreatic islets before detectable insulitis [11, 15], and blocking CXCL10 delays immune-mediated diabetes in these mice [16, 17]. Expressing CXCL10 in β-cells led to impaired β-cell function and spontaneous infiltration of lymphocytes into pancreatic islets [18]. Additionally, CXCL10 originating from β-cells was also shown to suppress β-cell proliferation [11]. Thus, mitigating signaling cascades that escalate the production of CXCL10 can be an effective strategy to protect β-cells, delay T1D onset and prevent further disease progression.

One therapeutic strategy could be to modulate specific SNARE exocytosis protein-regulated pathway(s) to protect β-cells from cytokine-mediated dysfunction and demise. Previously, β-cell-specific enrichment of the exocytosis protein, Syntaxin 4 (STX4), blocked proinflammatory cytokine-induced expression of chemokine ligands (i.e., CXCL9, CXCL10 and CXCL11) in β-cells, and significantly deterred conversion to autoimmune-diabetes in NOD mice [19, 20]. The STX4-regulatory protein, Double C-2 like domain beta (DOC2b), when enriched in β-cells of primary human islets, mouse islets, and β-cell lines, promoted their function and survival [21–24], and in vivo, countered the effects of multiple low dose streptozotocin [25] (MLD-STZ, a chemical-inducer of β-cell destruction in mice [26]). DOC2b, a 46–50 kDa protein, is composed of an N-terminal Munc13-interacting domain (MID) and C-terminal tandem Ca^2+^ and phospholipid-binding C2 domains (C2A and C2B). C2A and C2B domains of DOC2b transiently interact with Munc18a and Munc18c, respectively, and serve as a scaffolding platform to facilitate SNARE complex formation and glucose-stimulated insulin secretion (GSIS) [27, 28]. Also, DOC2b’s Ca^2+^ binding C2 domain is important for promoting GSIS in β-cells [25, 29].

T1D human islets harbor reduced levels of several exocytosis proteins, proteins largely known to control insulin secretory function, but to date, only the t-SNARE protein STX4 [30] and its regulator DOC2b [31], are implicated in protecting against T1D stressor-induced β-cell death in vivo. In particular, DOC2b protein levels in T1D human islet β-cells are nearly undetectable [32]. By contrast, DOC2b-enrichment in β-cells protects them from proinflammatory cytokine-induced endoplasmic reticulum (ER)-stress and apoptosis, preserving β-cell mass [25]. Even though this evidence demonstrated the beneficial contribution of DOC2b in preventing stress-induced β-cell destruction [25], the mechanistic details remain to be elucidated.

In the current study, we hypothesized that DOC2b enrichment mitigated cytokine-induced β-cell apoptosis via disrupting proinflammatory signaling cascades, attenuating CXCL10 signaling. Here, we tested this hypothesis using our global DOC2b heterozygous knockout mice (DOC2b^+/-^), DOC2b-enriched or-depleted primary human islets, and the rat INS-1/832/13 β-cell line challenged with T1D-related stressors.

## 2. MATERIALS AND METHODS

### 2.1. Human islets

Cadaveric non-diabetic pancreatic islets were obtained from the Integrated Islet Distribution Program and City of Hope Islet Cell Resource Center (Duarte, CA) and cultured as previously described [19, 24]. The donors (male and female) were between 20-57 years old (Suppl. Table 1).

### 2.2. Animals, multiple-low-dose streptozotocin (MLD-STZ) treatment, and mouse islet isolation

The Animal Use Protocol #23042 was approved by the Institutional Animal Care and Use Committee at the City of Hope (Duarte, CA). Eight-week-old male DOC2b^+/-^ [23] and wild-type (WT) mice were subjected to MLD-STZ treatment as before [25]. Subsequently, pancreata were collected and processed for immunohistochemistry studies. For cytokine treatment studies pancreatic mouse islets were isolated from 10- to 12-week-old female DOC2b^+/-^ and WT mice and cultured overnight, as previously described [33].

### 2.3. Immuno-confocal microscopy

For Immunohistochemistry, paraffin-embedded pancreatic tissue sections from mice were prepared and assessed as previously described [19, 25, 34]. Pancreatic sections were stained with guinea pig anti-insulin (1:50, Invitrogen), anti-NF-κB p65 (1:100, Cell Signaling Tech.), and rabbit anti-CXCL10 (1:100, Invitrogen) primary antibodies at 4°C overnight, and Invitrogen Alexa Fluor 488 (anti-guinea pig 1:100)-, 647(anti-rabbit, 1:300)-conjugated secondary antibodies for 1 h at room temperature. Nuclei were stained using 49,6-diamidino-2-phenylindole (DAPI) and were subsequently overlaid with Vectashield (Vectashield; Vector Laboratories, Burlingame, CA) mounting medium for imaging with a Zeiss 900 confocal microscope with a 40X objective.

### 2.4. RNA isolation, quantitative real-time PCR, siRNA-mediated knockdown of DOC2b

RNA isolation and quantitative real-time PCR were performed as before [19, 20]. Sequences for the primers used to detect CXCL10, CXCL9, STAT-1, ATF4, CHOP, and DOC2b genes are provided in Suppl. Table 2. Gene expression levels were normalized to GAPDH expression. For knockdown of DOC2b in INS-1 832/13 β-cells, the small-interfering (si) RNA oligonucleotides (50 nmol/L rat DOC2b, and control; ON-TARGET plus, Thermo Dharmacon, Lafayette, CO) were transfected into INS-1 832/13 β-cells using RNAiMax, and assessments were made 48 h later.

### 2.5. Immunoblotting

Total proteins were harvested using RIPA lysis buffer (Thermo Fisher Scientific, USA). Cleared detergent lysates were resolved by 10% or 15% (CXCL10, Cl. Caspase 3 blots) SDS-PAGE and transferred to polyvinylidene fluoride membranes for immunoblotting. Detailed information about the primary antibodies is provided (Suppl. Table 3). Goat anti-mouse and anti-rabbit horseradish peroxidase secondary antibodies (Bio-Rad Laboratories, Hercules, CA) were used at 1:5,000. Immunoreactive bands were visualized with Enhanced chemiluminescence (ECL), ECL Prime, or ECL Super-Signal reagents (GE Healthcare, USA) and imaged using the ChemiDoc gel documentation system (Bio-Rad Laboratories, Hercules, CA).

### 2.6. Cell culture, adenoviral transduction, cytokines, and cyclopiazonic acid (CPA) treatment

INS-1 832/13 β-cells were kindly provided by Dr. Christopher Newgard (Duke University Medical Center, Durham, NC) and were cultured as before [35]. The adenoviruses Ad.GFP, Ad. (rat) rDOC2b-GFP, and Ad. (Human) hDOC2B-MYCDDK were generated by ViraQuest (North Liberty, IA), as described before [24]. The Ad. Empty vector control for Ad. hDOC2b-MYCDDK was purchased from Viraquest, Inc. (North Liberty, IA). INS-1 832/13 β-cells between 55-65 passages were used for experiments. Human islets and rat INS-1 832/13 β-cells were transduced adenovirally as before [19, 20, 24]. For proinflammatory cytokine treatment, cells were cultured in a medium containing human (human islets) or rat (INS-1 832/13 β-cells) or mouse (mouse islets)-specific cytokine cocktail (100 ng/mL interferon-γ [IFN-γ], 10 ng/mL tumor necrosis factor-α [TNF-α], and 5 ng/mL interleukin-1β [IL-1β] (ProSpec-Tany Techno Gene, East Brunswick, NJ) [19, 20] for 1h (acute treatment) or ≥8h (chronic treatment). For concomitant CPA and cytokine treatments, INS-1 832/13 β-cells were pre-treated with 6.25 μM CPA (Sigma-Aldrich) or vehicle for 6h followed by exposure to cytokines for 1h. Post-treatment cleared detergent lysates were prepared using commercially available RIPA buffer for immunoblotting, or 1% NP40 lysis buffer (co-immunoprecipitation); RNA was isolated for quantitative real-time PCR.

### 2.7. Co-immunoprecipitation

Detergent (1% NP40)-solubilized protein lysates (500 μg)/purified recombinant proteins were co-immunoprecipitated using GFP-Trap Magnetic Agarose/DDK-Fab-agarose beads, respectively (Proteintech, Rosemont, IL) per the manufacturer’s instructions.

### 2.8. Subcellular Fractionation of INS-1 832/13 β-cells

Transduced INS-1 832/13 β-cells were treated with a proinflammatory cytokine cocktail for ≥8h and were partitioned into nuclear and non-nuclear fractions using NE-PER Nuclear and Cytoplasmic Extraction Kit (Thermo Fisher Scientific, USA). Sub-cellular fractions were used for immunoblotting analyses to detect T-STAT-1.

### 2.9. Statistical Analysis

Results were evaluated for statistical significance using an unpaired two-tailed Student t-test for comparison of two groups or one-way ANOVA for ≥ two groups using Prism version 10.0 (GraphPad Software, La Jolla, CA). Data are expressed as the mean ± SEM.

### 2.10. Data and Resource Availability

The data sets generated during and/or analyzed during the current study are available from the corresponding author upon reasonable request. No applicable resources were generated during this study.

## 3. RESULTS

### 3.1. DOC2b depletion exacerbates stress induced CXCL10 expression in primary mouse islets and INS-1 832/13 β-cells

Previously, we showed that DOC2b^+/-^ mice were highly susceptible to MLD-STZ-induced glucose dyshomeostasis and β-cell apoptosis [25] and that DOC2b protected against cytokine-induced apoptosis [25]. Further, CXCL10 is known to drive β-cell death in human T1D, and autoimmune diabetes in mice [11, 13, 14]. Thus, to determine if DOC2b deficiency confers β-cell susceptibility to stress-induced CXCL10 expression, we evaluated CXCL10 levels in MLD-STZ challenged DOC2b^+/-^ and WT mouse islets. Islet CXCL10 levels were substantially elevated in DOC2b^+/-^ versus WT islets, nearly 2-fold higher specifically in the insulin-stained islet area (Fig. 1A, bar graph). In the absence of MLD-STZ challenge, DOC2b^+/-^ islets had minimal CXCL10 levels (Fig. 1A). Next, we challenged DOC2b^+/-^ and WT primary islets from 10-12-week-old female mice with a proinflammatory cytokine cocktail that mimics the T1D pancreatic milieu [36] to determine if DOC2b deficiency increases susceptibility to proinflammatory cytokine-induced CXCL10 expression. Chronic exposure (C, ≥ 8 h) to a cocktail of cytokines was used to simulate T1D stress on DOC2b^+/-^ islets, which showed a significant escalation in CXCL10 mRNA levels, versus that of WT islets (Fig. 1B). These results suggested inverse correlation of DOC2b and cytokine-induced CXCL10 protein levels in mouse islets. Consistent with this, endogenous DOC2b knockdown (siDOC2b) in rat INS-1 832/13 β-cells challenged with the cytokine cocktail resulted in a ∼1.5-fold-increase in CXCL10 mRNA levels versus siControl (Fig. 1C-E), within 8 h of cytokine exposure. The results suggest that DOC2b deficiency increases β-cell susceptibility to cytokine-induced CXCL10 expression in vivo and in vitro.

**FIGURE 1.**
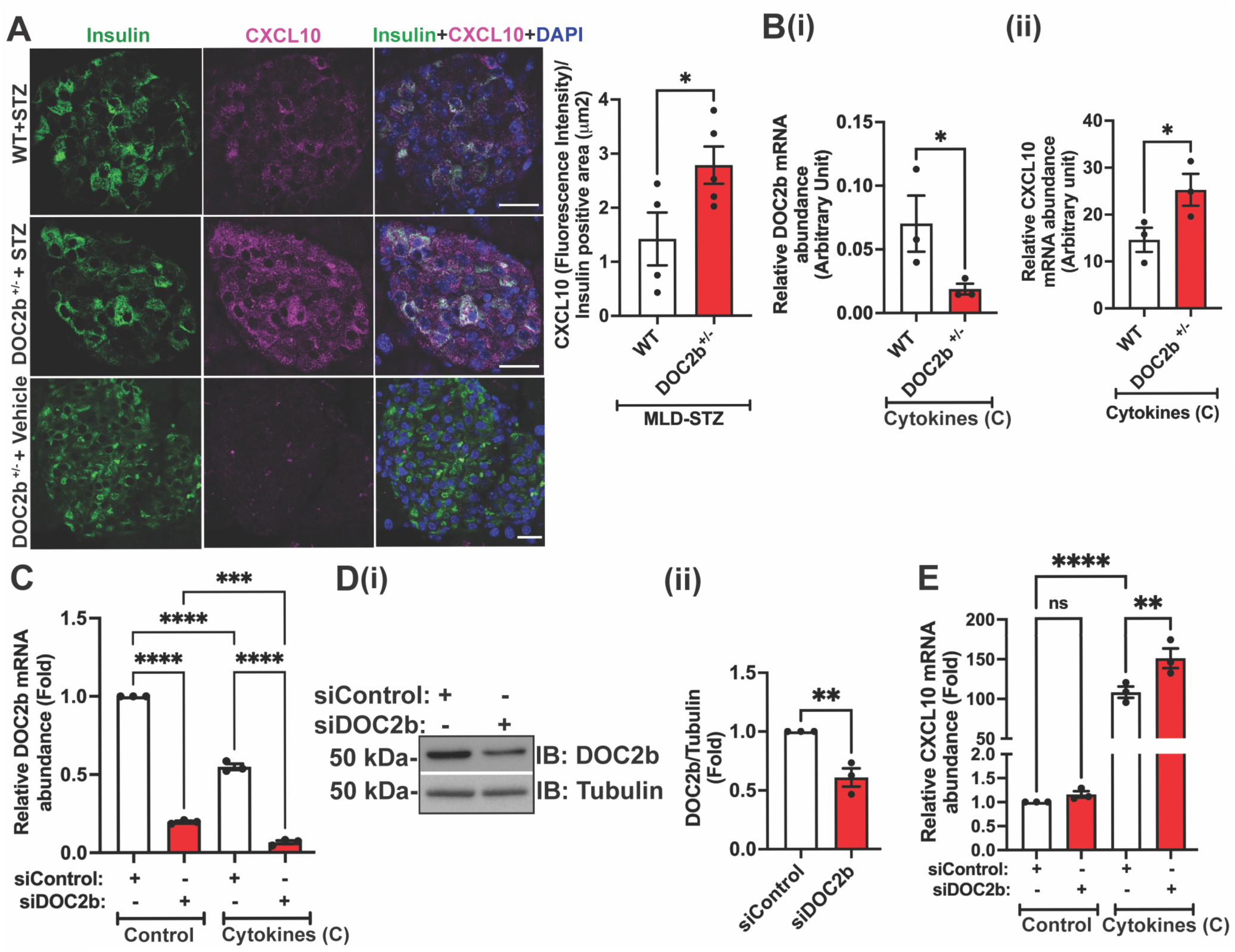
DOC2B is required for attenuating stress-induced CXCL10 expression in β-cells. (A) Insulin (green) and CXCL10 (purple) immunofluorescent staining; bar graph quantitation of CXCL10 intensity in insulin+ cells in mouse pancreata, representative of n = 4-5 mice/group. Bar = 20 μm. DAPI (blue), nuclear staining. (B) qPCR quantitation of relative DOC2b (i) and CXCL10 (ii) mRNA abundance in cytokine cocktail-exposed (chronic (C), ≥ 8h) mouse islets (n=3 mice/group). (C-E) Depletion of endogenous DOC2b using siRNA-mediated knockdown of rat DOC2b (siDOC2b) vs. scrambled control (siControl) in INS-1 832/13 β-cells is furthered by chronic exposure to a cytokine cocktail (n = 3 independent cell passages). (C) qPCR analysis of DOC2b mRNA levels. (D) Validation of DOC2b depletion by immunoblot (IB) analysis (i) and quantitation of DOC2b protein (ii). (E) qPCR quantitation of CXCL10 levels from the same cells as in (C-D). Bars represent the mean ± SEM. *P<0.05, **P< 0.01, ***P<0.001, ****P<0.0001, ns= not significant.

### 3.2. DOC2b overexpression attenuates CXCL10 expression in cytokine-challenged human islets and INS-1 832/13 β-cells

Next, we asked if DOC2b enrichment attenuates chemokine ligand expression in cytokine-challenged islets/β-cells. We used adenovirally transduced (DOC2b versus control empty vector) non-diabetic human primary islets challenged with a proinflammatory cytokine cocktail [19] to assess CXCL10 mRNA and protein levels. Compared to control, cytokine-induced CXCL10 gene and protein expressions were significantly attenuated (∼50%) in DOC2b-transduced human islets; the lack of detectable CXCL10 protein in the absence of cytokine treatment further confirmed cytokine-responsive CXCL10 protein expression in human islets (Fig. 2Ai-v). Given that human islets consist of cell types beyond β-cells, we evaluated CXCL10 expression in cytokine-responsive INS-1 832/13 β-cells transduced with either DOC2b-GFP or vector control (GFP); DOC2b-GFP overexpression resulted in a > 50% reduction in CXCL10 mRNA expression in cytokine-stressed β-cells versus control (Fig. 2Bi-iii). Consistent with this, we also observed that cytokine-induced CXCL9 expression was significantly reduced (∼50%) in DOC2b-overexpressing human islets and INS-1 832/13 β-cells (Suppl. Fig. 1). Vector-transduced human islets showed increased apoptosis via cleaved caspase 3 post-chronic cytokine challenge versus control validating cytokine action (Suppl. Fig. 2A); insulin content was not statistically different (Suppl. Fig. 2B) [37]. In cytokine-stressed vector transduced INS-1 832/13 β-cells, significant ablation of glucose-stimulated insulin secretion (GSIS; Suppl. Fig. 2C), insulin content (Suppl. Fig. 2D), and elevated apoptosis (Suppl. Fig. 2E) versus control were observed, validating cytokine action. DOC2b overexpression also significantly increased GSIS in chronic cytokine-stressed INS-1 832/13 β-cells vs vector control (Suppl. Information, Suppl. Fig. 3).

**FIGURE 2.**
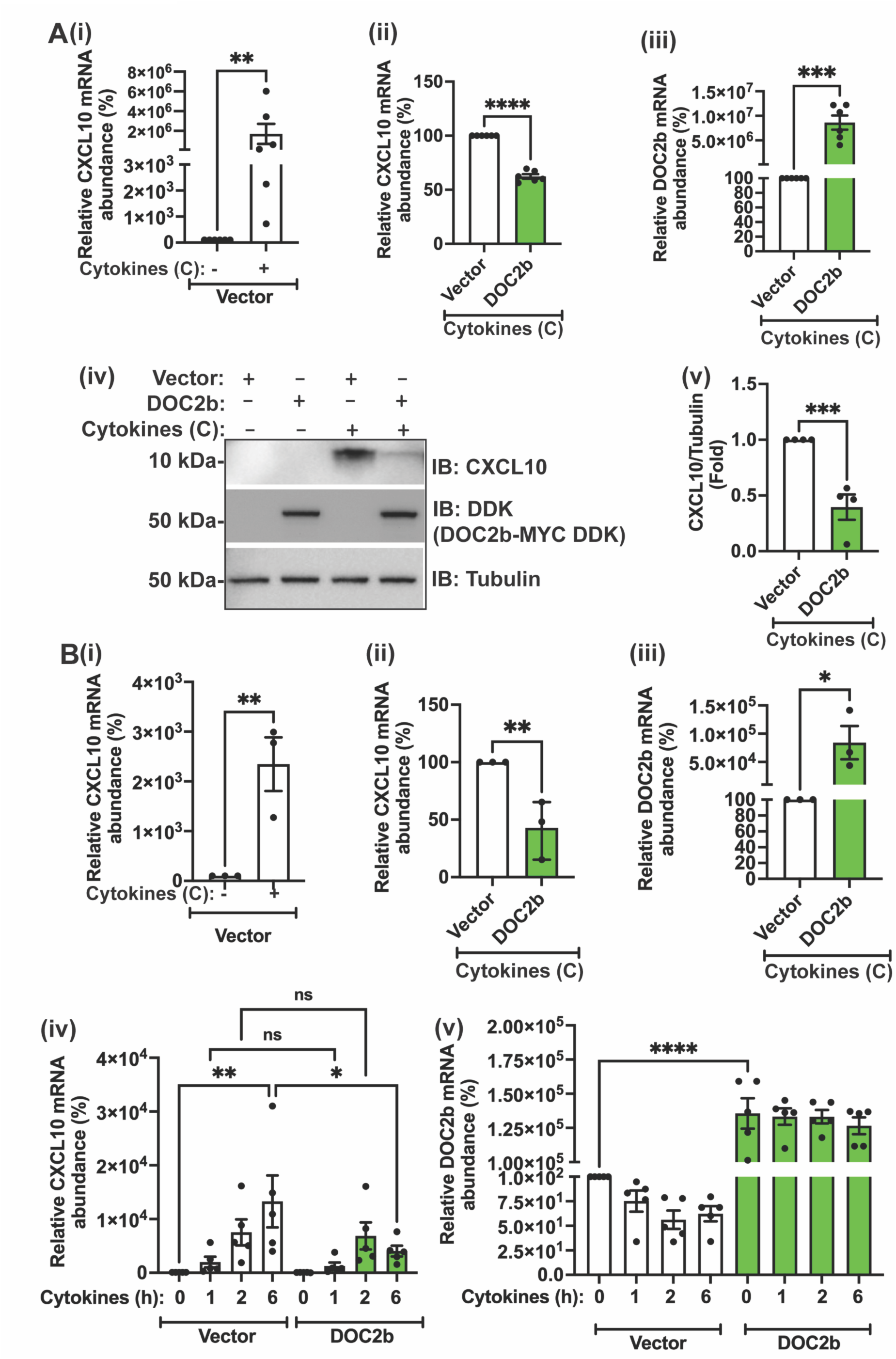
DOC2B overexpression protects against proinflammatory cytokine-induced CXCL10 gene expression in human islets and INS-1 832/13 β-cells. (A) DOC2b- or control (vector)-transduced non-diabetic human islets were exposed to a proinflammatory cytokine cocktail (chronic treatment (C), ≥ 8h). (i-iii) qPCR quantitation of CXCL10 (i-ii) and DOC2b (iii); N=6 donors/group. (iv-v) Immunoblot analysis (IB), representative of 4 independent human donor islet batches (iv), and quantitation of CXC10 protein levels (v). Tubulin was used as the loading control. (B) DOC2b-GFP or GFP-control (vector)-transduced INS-1 832/13 β-cells were exposed to a cytokine cocktail for varying durations. (i-iii) Chronic exposure: qPCR quantitation of CXCL10 (i-ii), and DOC2b (iii); N=3 independent cell passages. (iv-v) qPCR quantitation of CXCL10 (iv) and DOC2b (v) across an acute time course of exposure to cytokine cocktail; n=5 independent cell passages. (A-B) Bars represent the mean ± SEM *P<0.05, **P<0.01, ***P<0.001, ****P<0.0001, ns= not significant.

Next, to determine the temporal dynamics of DOC2b action on cytokine-induced CXCL10 gene expression, DOC2b-GFP- or vector control-transduced INS-1 832/13 β-cells were exposed to proinflammatory cytokine cocktail for up to 6 h. Within 6 h, attenuation of CXCL10 mRNA with DOC2b-GFP overexpression was observed (Fig. 2Biv); DOC2b enrichment was maintained during 6 h of cytokine challenge (Fig. 2Bv). These results indicate that DOC2b protects primary human islets and rat β-cells against cytokine-induced chemokine ligand expression.

### 3.3. DOC2b mitigates CXCL10 expression in human islets and INS-1 832/13β-cells in response to individual cytokines

To determine the contribution of DOC2b in mitigating CXCL10 gene expression in response to individual cytokines, DOC2b, or control vector-transduced non-diabetic primary human islets, were challenged with either IL-1β, TNF-α, or IFN-γ. As previously shown in human islets [38], IFN-γ induced the highest CXCL10 gene expression (> 20-30-fold) (Fig. 3Ai). Intriguingly, DOC2b overexpression in human islets significantly prevented the IL-1β-or IFN-γ-triggered increases in CXCL10 mRNA expression versus vector control (Fig. 3Ai-iii); DOC2b was without effect upon TNF-α-induced CXCL10 (Fig. 3Aiv). Comparable DOC2b expression levels between IL-1β-, TNF-α-, and IFN-γ-treated human islets did not account for the differential effects observed above (Fig. 3Av). Cytokine challenge is known to upregulate ER stress in primary islets and β-cell lines [39–42]. ER stress marker ATF4 mRNA levels were elevated in cytokine-challenged vector-transduced human islets (Suppl. Fig.4A).

**FIGURE 3.**
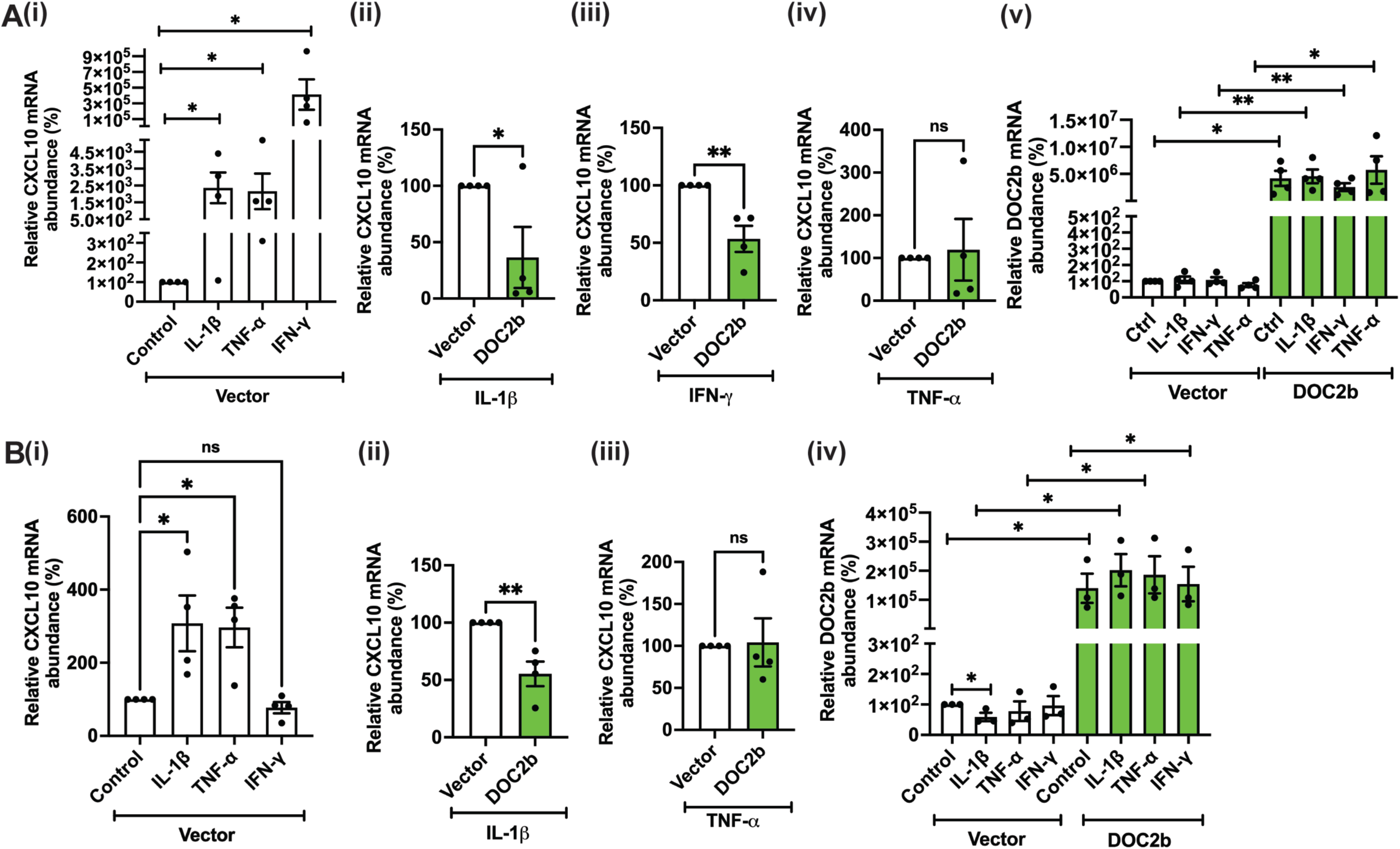
DOC2b overexpression protects against IL-1β and IFN-γ induced CXCL10 gene expression in human islets and IL-1β induced CXCL10 gene expression in INS-1 832/13 β- cells. (A) DOC2b- or control (vector)-transduced non-diabetic human islets were exposed to proinflammatory cytokines (chronic treatment, C; ≥ 8h). (i-v) qPCR quantitation of CXCL10 (i-iv), and DOC2b (v); n= 4 donors/ group. (B) DOC2b-GFP- or vector control (GFP)-transduced INS-1 832/13 β-cells were exposed to cytokine stress. (i-iv) qPCR quantitation of CXCL10 (i-iii), and DOC2b (iv); n= 3-4 independent passages. (A-B) Bars represent the mean ± SEM. *P<0.05, **P< 0.01, ns= not significant.

We next assessed for DOC2b-mediated CXCL10 gene downregulation in INS-1 832/13 β- cells. DOC2b-GFP- or control vector-transduced β-cells were challenged with either IL-1β, TNF-α, or IFN-γ. Increased CHOP expression (an ER stress marker) in the vector-transduced INS-1 832/13 β-cells versus control confirmed the effectiveness of cytokine challenge (Suppl. Fig.4B). Like human islets, CXCL10 upregulation was observed with IL-1β or TNF-α-treatment in GFP-transduced β-cells (Fig. 3Bi). However, unlike human islets, IFN-γ-treated INS-1 832/13 β-cells failed to induce CXCL10 expression (Fig. 3Bi), as previously published [9]. DOC2b-enrichment only reduced IL-1β-induced CXCL10 gene expression and lacked effect upon TNF-α-induced CXCL10 (Fig. 3Bii-iv), like that seen in human islets (Fig. 3Aiv), despite comparable DOC2b mRNA levels in IL-1β- and TNF-α-treated INS-1 832/13 β-cells (Fig. 3Biv). Altogether, these results suggest that DOC2b enrichment mitigates IL-1β-induced CXCL10 expression in both human islets and INS-1 832/13 β-cell line, and IFN-γ-induced CXCL10 expression in human islets.

### 3.4. DOC2b levels inversely correlate with NF-κB p65 levels in stressed islets

NF-κB p65 is a regulator of CXCL10 in β-cells [9] and is induced in STZ-challenged mouse islets [43]. Thus, total NF-κB p65 levels in MLD-STZ challenged DOC2b^+/-^ and WT mouse islets were examined. MLD-STZ challenged DOC2b^+/-^ islets showed elevated total (T) NF-κB p65 levels versus WT (Fig. 4Ai-ii) and unchallenged DOC2b^+/-^ islets (Fig. 4Ai). Furthermore, chronically cytokine-stressed DOC2b-enriched human islets had reduced total NF-κB p65 protein levels versus control (Fig. 4Bi-ii). Taken together, DOC2b levels inversely correlated with total NF-κB p65 levels in stressed islets, suggesting a link between DOC2b and inflammatory signaling.

**FIGURE 4.**
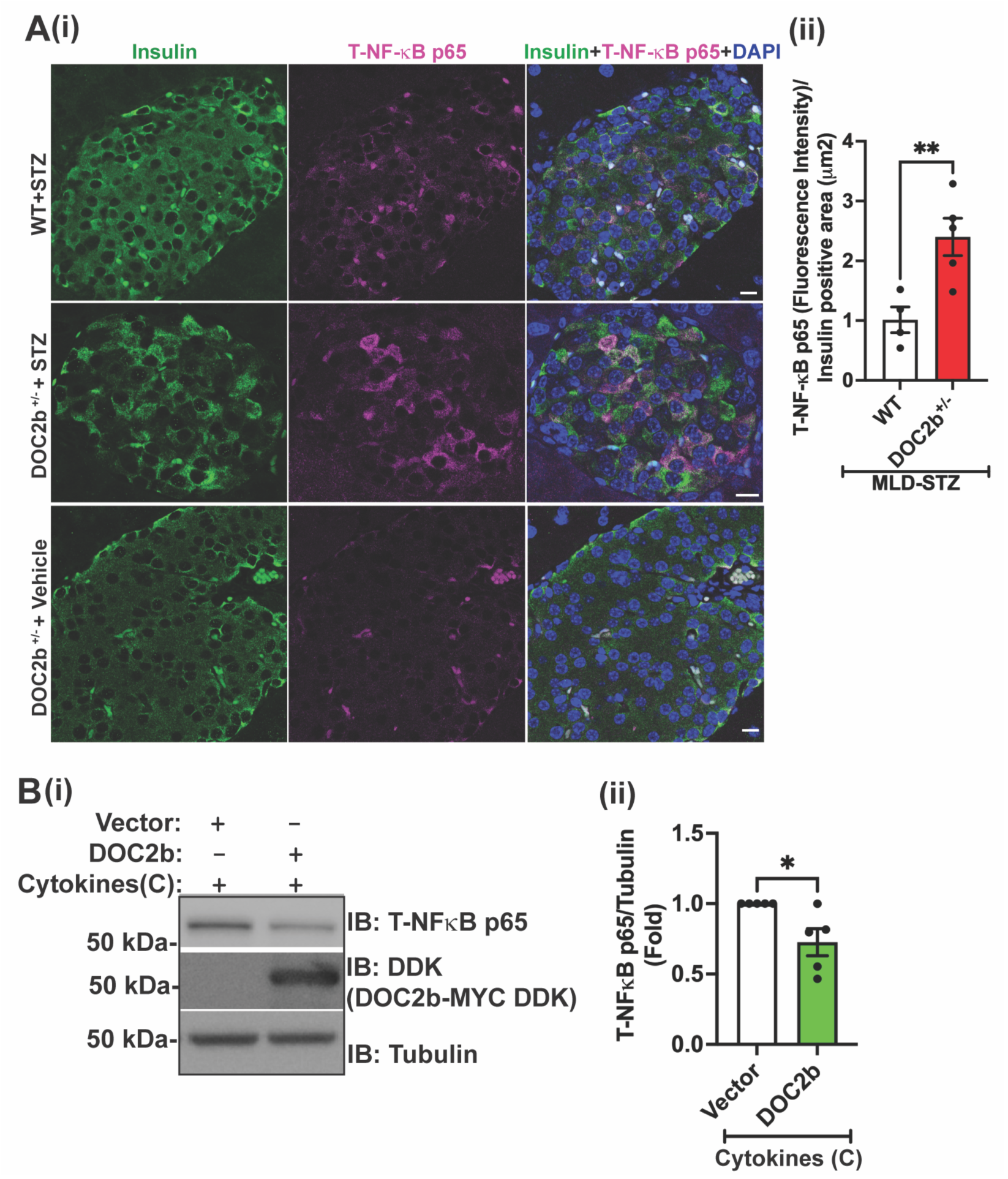
DOC2b levels are inversely proportional to total NF-κB p65 protein levels in stressed mouse and human islets. (Ai-ii) Insulin (green) and total NF-κB p65 (purple) immunofluorescent staining (i); quantitation of NF-κB p65 intensity in insulin+ cells in mouse pancreata (n= 4-5 mice/group). Bar = 20 μm. DAPI (blue), nuclear staining (ii). (B) DOC2b- or control (vector)-transduced non-diabetic human islets were treated with a proinflammatory cytokine cocktail (chronic treatment (C), ≥ 8h). (i-ii) Immunoblot analysis (IB), representative of 5 independent human donor islet batches (i), and quantitation of total (T) NF-κB p65 protein levels (ii). Tubulin was used as the loading control. Bars represent the mean ± SEM. *P<0.05.

### 3.5. DOC2b associates with NF-κB p65 pathway and STAT-1 in cytokine-challenged INS-1 832/13 β-cells

Assessment of our prior mass spectrometry analysis [24] revealed multiple putative inflammatory/apoptosis-related proteins (e.g., IKKβ, IKKα, STAT-1) affiliating with DOC2b-GFP versus control (Suppl. Table 4). In cytokine-stressed β-cells, NF-κB p65 and STAT-1 regulate CXCL10 expression [9, 44]. As such, we determined if DOC2b associates with IKKβ, NF-κB p65- pathway factors, and/or STAT-1, in cytokine-stressed β-cells at a time shown to precede CXCL10 induction [20]. In GFP/DOC2b-GFP-transduced INS-1 832/13 β-cells cytokine stressed for 1h (Acute [A]), DOC2b-GFP associated with NF-κB p65, IKKβ, and STAT-1 (Fig. 5A). Further, DOC2b coimmunoprecipitated IKKβ/STAT-1 proteins in a cell-free system, using all recombinantly expressed and purified DOC2b, IKKβ-MYCDDK, and STAT-1-MYCDDK proteins, suggestive that these proteins may be capable of protein-protein interactions (Fig. 5B, C). AlphaFold analysis also predicts the interactions of DOC2b’s C-terminal C2B and N-terminal domains with IKKs, while the C2B domain alone was predicted to interact with NF-κB p65 and STAT-1 (Suppl. Information, Suppl. Fig. 5A-G). Interestingly, DOC2b’s Y301 formed part of the interface with NF-κB p65, but not with STAT-1 (Suppl. Information, Suppl. Fig. 5G-H). Functionally, both DOC2b-^WT^ and ^Y301F^ overexpression attenuated CXCL10 expression (Suppl. Fig. 6i-ii) in cytokine-challenged INS-1 832/13 β-cells, whilst the DOC2b tandem C2AB domain failed to attenuate CXCL10 mRNA abundance (Suppl. Fig. 6iii-iv). These results support the concept of DOC2b’s regulation of CXCL10 via IKKβ, NF-κB p65, and STAT-1 pathways.

**FIGURE 5.**
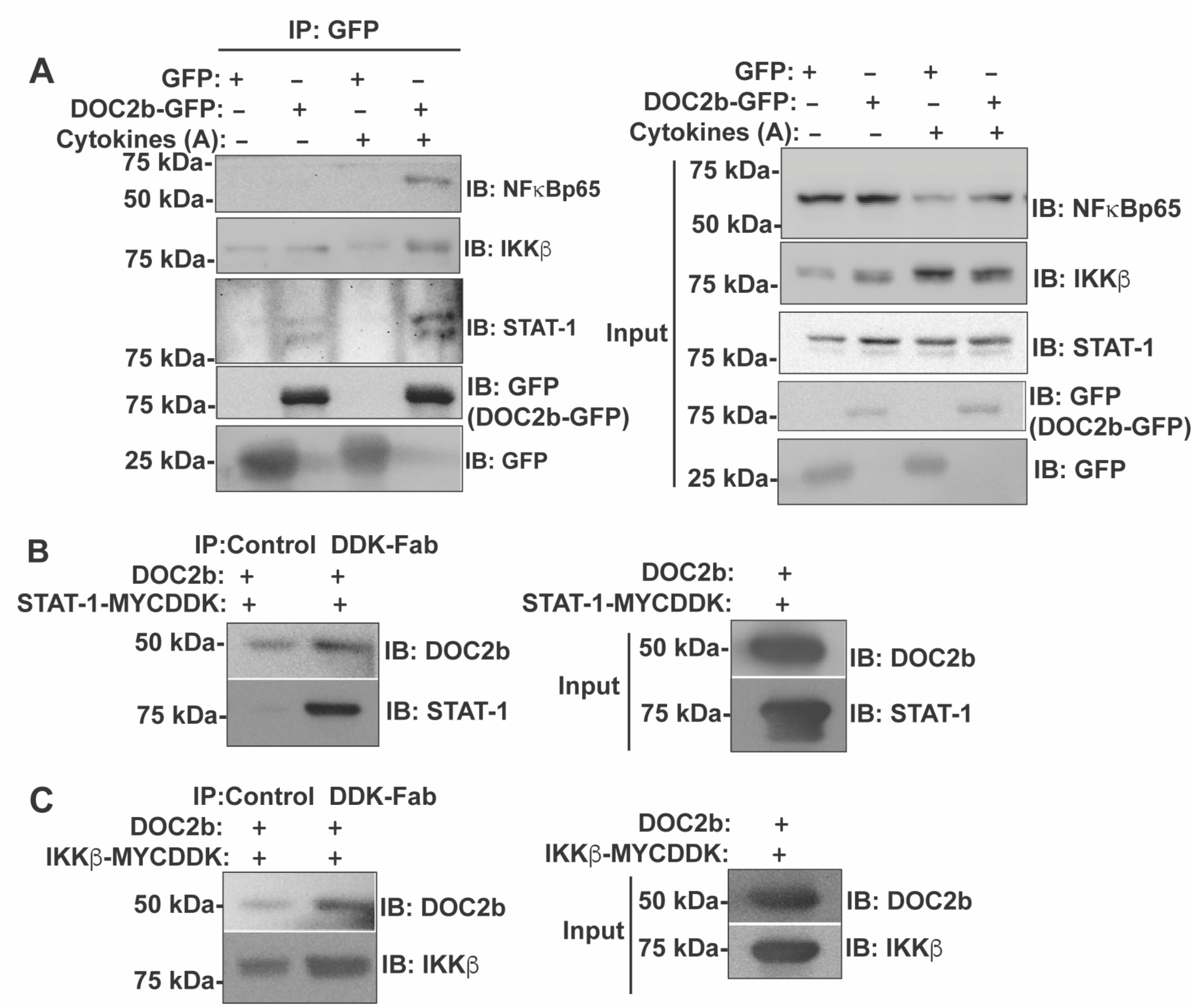
DOC2b associates with NF-κB p65, IKKβ and STAT-1 proteins in cytokine challenged INS-1 832/13 β-cells. (A) INS-1 832/13 β-cells were transduced with either GFP control/DOC2b-GFP adenoviruses and were exposed acutely to the pro-inflammatory cytokine cocktail (rat IL1β+TNFα+IFN-γ) (1h acute treatment, A). Immunoblot (IB) detection of co-immunoprecipitated (IP) proteins under stated conditions; images represent n=3 independent experiments. (B-C) Purified recombinant proteins (DOC2b and IKKβ-MYCDDK/STAT-1-MYCDDK) were combined in equimolar amounts for subsequent immunoprecipitation (IP) with Anti-DDK-Fab-agarose beads, and precipitated proteins were resolved by 10% SDS-PAGE for subsequent immunoblot detection (IB). IP with control beads was used as a negative control for non-specific binding. Input proteins were sampled from reactions to confirm the presence of DOC2b, IKKβ-MYCDDK, and STAT-1-MYCDDK. Equivalent volumes captured from the reaction were used to confirm the input of each protein in the reaction. Images represent n=3 independent experiments.

### 3.6. DOC2b enrichment attenuates IKKβ levels in cytokine-challenged human islets and INS-1 832/13 β-cells

We next determined if DOC2b overexpression altered the cytokine-induced activation of upstream canonical NF-κB signaling factors in human islets and INS-1 832/13 β-cells. In β-cells, cytokine-induced CXCL10 expression is regulated by NF-κB p65 (canonical NF-κB pathway factor) and STAT-1 signaling [9, 44]. The cytokine-induced activation of the canonical NF-κB pathway is mediated by the upstream inhibitor of κB (IκB) kinase (IKK) complex [45, 46]. The subunits IKKβ and IKKα of the IKK complex are associated as homodimers or heterodimers and are bound to the regulatory subunit, IKKγ [47]. In non-stimulated conditions, the NF-κB inhibitory proteins, IκBβ and IκBα are bound with NF-κB in the cytoplasm, inhibiting its nuclear transport. Following cytokine stimulation activated IKK complex leads to phosphorylation of IκBα, IκBβ, followed by their proteasome-mediated degradation, thus releasing NF-κB members to form dimers (p50 and p65, homo or heterodimers). Following this, NF-κB dimers translocate to the nucleus and regulate gene expression [45, 46].

To understand the link between DOC2b and canonical NF-κB signaling DOC2b-transduced non-diabetic primary human islets were exposed to an acute (A, 1 h) cytokine cocktail to enable detection of early events of canonical NF-κB signaling [20]: the activated-and total protein levels of IKKβ and IKKα were assessed. No changes were observed in total IKKβ protein level between control and DOC2b-enriched human islets in the absence of cytokines (Suppl. Fig. 7). Cytokine-mediated activation of IKK (p-IKKβ^serine 180^) was observed in vector-transduced control islets (Suppl. Fig. 8A). DOC2b enrichment significantly reduced both cytokine-induced total and activated levels of IKKβ (Fig. 6A-B); the activated/total ratio of IKKβ was not altered (Fig. 6A-B). Further, only minimal changes were observed in cells with DOC2b enrichment upon total IKKα level (Fig. 6A-B) or IKKα activation (p-IKKα^serine 176^), as compared with control cells (Suppl. Fig. 8A).

**Figure 6.**
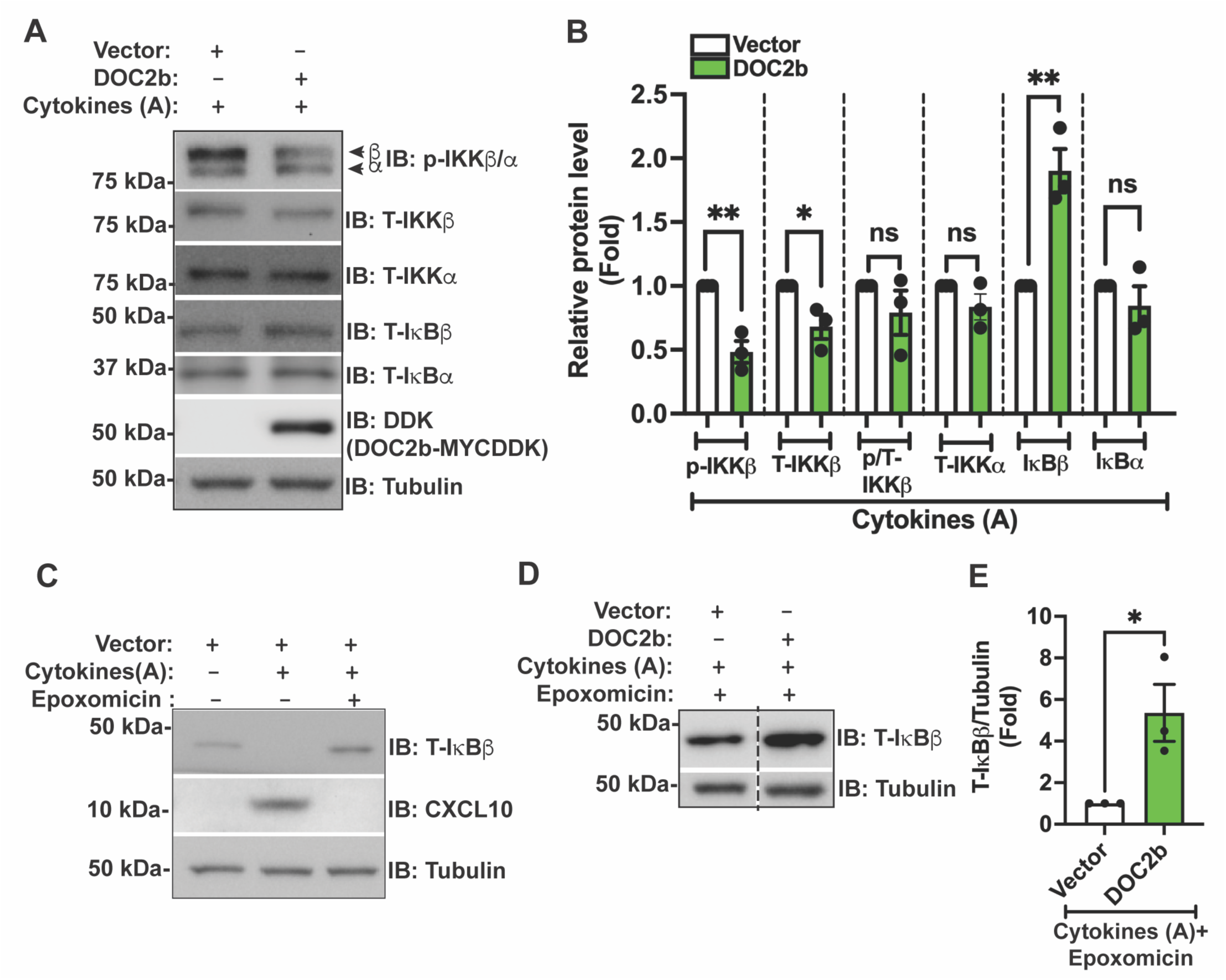
DOC2b enrichment reduces activated and total IKKβ protein and increases total IκBβ protein levels in human islets following acute cytokine treatment. (A-B) DOC2b- or control (vector)-transduced non-diabetic human islets were exposed to a proinflammatory cytokine cocktail (acute treatment (A), 1h). (A-B) Immunoblot analysis (IB) representative of 3 independent human donor islet batches (A), and quantitation of protein levels (B). Tubulin was used as the loading control. (C-E) DOC2b- or vector (control)-transduced non-diabetic human islets were treated with a proinflammatory cytokine cocktail (acute treatment (A), 1h) and proteasome inhibitor epoxomicin (10μM); n=3 independent donors. For epoxomicin treatment, cells were pre-incubated with epoxomicin for 30 min, followed by incubation with epoxomicin plus a proinflammatory cytokine cocktail for an additional 1 h. Immunoblot analysis (IB) (C-D), and quantitation of total (T) IκBβ protein levels (E). Tubulin was used as the loading control. Bars represent mean ± SEM. *P<0.05, **P< 0.01, ns= not significant. A vertical dashed line indicates the splicing of lanes from within the same gel exposure.

These findings for total and p-IKKβ were recapitulated in cytokine-challenged DOC2b- enriched INS-1 832/13 β-cells (Suppl. Fig. 9A-B); p-IKKα detection was low and largely unresponsive to the acute cytokine challenge (Suppl. Figs. 8B, 9A-B). As with DOC2b-enriched human islets, activated/total IKKβ ratio was not altered in DOC2b-enriched INS-1 832/13 β-cells (Suppl. Figs. 9A-B).

### 3.7. DOC2b enrichment attenuates cytokine-induced IκBβ proteasomal degradation in human islets

Next, to seek out if DOC2b overexpression alters IKK downstream signaling in cytokine-stressed human islets, we assessed IκBβ and IκBα levels. Based on the observed mitigation of upstream p-IKKβ and T-IKKβ protein levels in DOC2b-enriched human islets (Fig. 6A-B), we evaluated DOC2b impact upon downstream signaling elements of IKK: IκBβ and IκBα. DOC2b-enriched human islets challenged with cytokines selectively contained ∼ 2-fold higher IκBβ, but not IκBα (Fig. 6A-B). Per the known role of proteasome in degradation of activated (IKK mediated phosphorylation)-IκBβ in β-cells [20], we tested the effects of proteasome inhibition (with epoxomicin, 10 μM [48]). Epoxomicin was validated to ablate the cytokine-induced loss of IκBβ, and induction of downstream CXCL10 expression, in vector-transduced human islets (Fig. 6C), and further enhanced IκBβ protein levels in islets enriched for DOC2b, to levels ∼5-fold higher than matched control islets (Fig. 6D-E). Complete inhibition of CXCL10 with concomitant cytokine and epoxomicin treatment in vector-transduced islets prevented the investigation of DOC2b-specific action on CXCL10 expression under these conditions (Fig. 6C). Altogether, these results indicated that DOC2b-inhibition of IKKβ-IκBβ signaling in human islets counteracts cytokine-induced IκBβ degradation by proteasome machinery in human islets.

### 3.8. DOC2b overexpression attenuates STAT-1 expression in cytokine-challenged human islets and INS-1 832/13 β-cells

p-STAT-1^Tyr701^ partially regulates IL1β+IFN-γ-induced CXCL10 expression in β-cells [9, 44]. Based on DOC2b’s mitigating action against IFN-γ-induced CXCL10 production in human islets (Figs. 2-3), and DOC2b-STAT-1 association in β-cells (Fig. 5, Suppl. Table 4, Suppl. Fig. 5H), we next investigated if DOC2b overexpression alters cytokine-induced STAT-1 signaling in human islets. Indeed, DOC2b overexpression significantly reduced activated (p-STAT-1^Tyr701^) and total STAT-1 levels in human islets following the acute cytokine cocktail challenge (Fig. 7Ai-ii). Marked increase of total STAT-1 mRNA and protein in control (vector) transduced human islets following chronic (≥ 8h) cytokine challenge confirmed cytokine-responsive STAT-1 expression in human islets (Fig. 7B, Ci). Again, compared to the control, chronic-cytokine-induced STAT-1 mRNA and protein levels were attenuated (∼40%) in DOC2b-transduced human islets (Fig. 7B-C). Similar to IKKβ, DOC2b enrichment in unstressed human islets did not impact STAT-1 levels (Fig. 7Ci, Suppl. Fig. 7), suggesting DOC2b is a negative regulator of cytokine-induced STAT-1 in islets.

**Figure 7.**
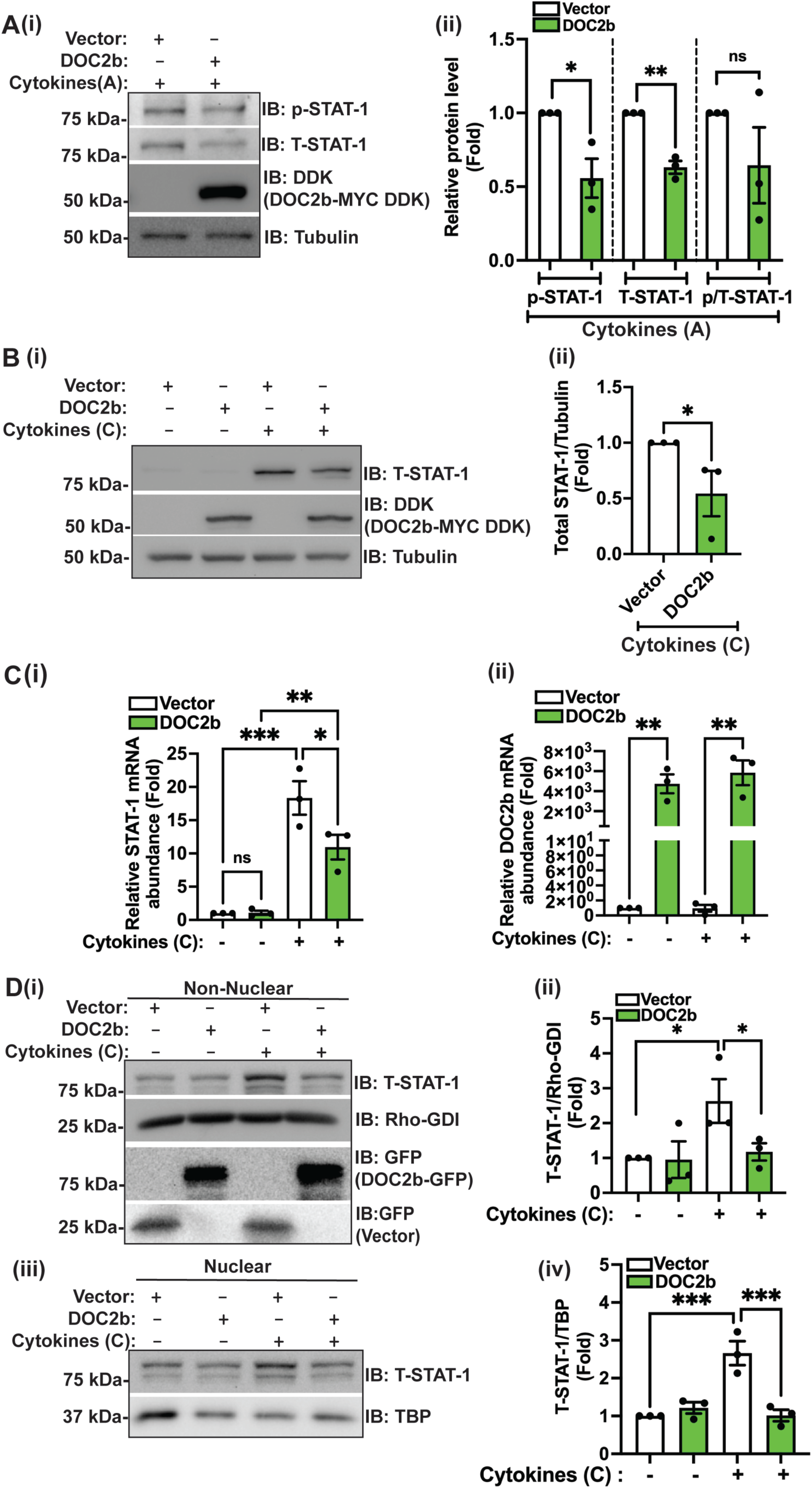
DOC2b enrichment prevents cytokine-induced STAT-1 protein escalation in human islets and INS-1 832/13 β-cells. (A) DOC2b- or control (vector)-transduced non-diabetic human islets were treated with a proinflammatory cytokine cocktail (acute treatment (A), 1h). (i-ii) Immunoblot analysis (IB) representative of 3 independent human donor islet batches (i), and quantitation of the indicated proteins (ii). Tubulin was used as the loading control. (B) DOC2b- or control (vector)-transduced non-diabetic human islets were treated with a proinflammatory cytokine cocktail (chronic treatment (C), ≥ 8h). Quantitation of STAT-1 (i) and DOC2b (ii); n=3 independent sets of donor islets. (D) DOC2b-GFP- or vector control (GFP)-transduced INS-1 832/13 β-cells were treated with a proinflammatory cytokine cocktail (chronic treatment (C), ≥ 8h) followed by subcellular fractionation into nuclear and non-nuclear fractions. (i, iii) Immunoblot analysis representative of 3 independent cell passages, and (ii, iv) quantitation of protein levels for the indicated proteins. Rho-GDI and TATA Binding Protein (TBP) were used as purity markers of non-nuclear and nuclear fractions, respectively. Bars represent mean ± SEM. *P<0.05, **P< 0.01, ***P<0.001, ns= not significant.

As with DOC2b-enriched human islets, total STAT-1 level was also reduced in DOC2b- enriched INS-1 832/13 β-cells following acute cytokine exposure (Suppl. Fig. 9A, C). Activated STAT-1 is known to translocate into the nucleus where it regulates inflammatory gene expression [49]. Non-nuclear and nuclear fractions prepared from GFP-transduced INS-1 832/13 β-cells showed a significant increase (3-4-fold) of T-STAT-1 protein following chronic cytokine-exposure (≥ 8h) versus unstressed control (Fig. 7D). This confirmed that cytokine-responsive STAT-1 protein expression changes in rat β-cell line. Consistent with DOC2b’s early reduction of T-STAT- 1 in human islets and β-cells (Fig. 7A, Suppl. Fig. 9A, C), both non-nuclear and nuclear fractions of DOC2b-GFP enriched INS-1 832/13 β-cells showed significant ∼ 50% reduction of chronic cytokine-induced STAT-1 protein level, as compared with the control (Fig. 7D). These results are consistent with the concept of DOC2b as an upstream regulator of cytokine-induced STAT-1 signaling in β-cells.

### 3.9. DOC2b overexpression modulates cytokine-induced IKKβ signaling and STAT-1 levels in mild-ER-stressed β-cells

Mild ER stress is known to sensitize rat β-cells to cytokine-induced inflammation and apoptosis [40, 41], and ER stress is a known regulator of STAT-1 signaling [50]. Further, in cytokine-mediated β-cell inflammation and apoptosis, there is evidence for crosstalk between ER-stress and NF-κB pathways [40, 41, 51]. We previously revealed that DOC2b can mitigate cytokine-induced ER stress and apoptosis in β-cells [25]. Thus, we asked if DOC2b overexpression can modulate cytokine-induced IKKβ signaling and STAT-1 levels in mild-ER-stressed β-cells. GFP/DOC2b-GFP-transduced INS-1 832/13 β-cells were pre-treated with low-dose ER-stress inducer CPA [40, 41], subsequently acutely exposed to cytokines, and IKKβ and STAT-1 levels assessed. Increased p-eIF2α/T-eIF2α level in GFP-CPA+Cytokines (Fig. 8A) vs GFP-control cells confirmed upregulation of ER-stress. DOC2b-GFP-CPA+Cytokines vs GFP-CPA+Cytokines β-cells showed significant attenuations of p-eIF2α/T-eIF2α level (Fig. 8A, B), p-and T-IKKβ (Fig. 8A, C-D), T-STAT-1 (Fig. 8A, E); p/T- IKKβ ratio was unchanged (Fig. 8A, F). Intriguingly, DOC2b markedly reduced the T-IKKβ level to below basal (the level seen without cytokine+CPA exposure of the GFP-control cells) (Fig. 8A, D). These results suggest that DOC2b overexpression can modulate cytokine-induced IKKβ signaling and STAT-1 levels in mild-ER-stressed β-cells.

**Figure 8.**
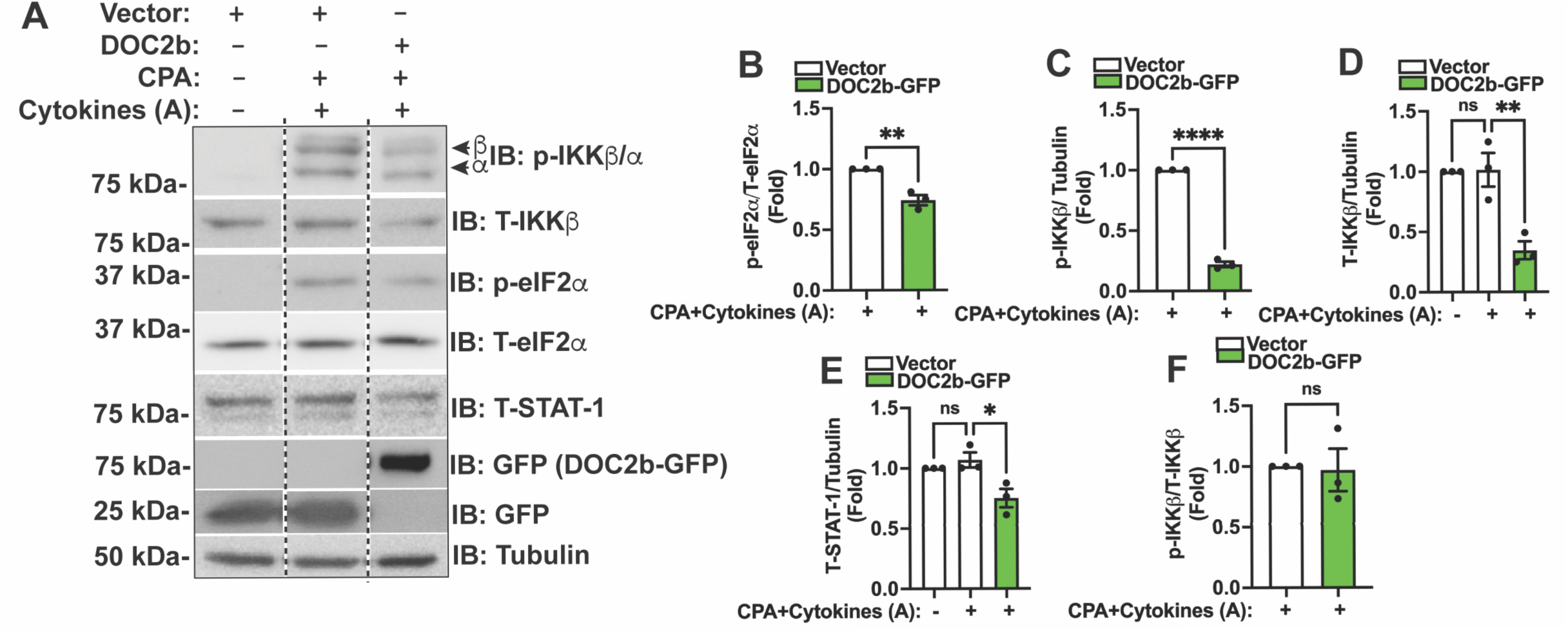
DOC2b overexpression modulates cytokine-induced IKKβ signaling and STAT-1 levels in mild-ER-stressed β-cells. DOC2b-GFP- or vector control (GFP)- transduced INS-1 832/13 β-cells were pre-treated with/without 6.26 μM CPA/vehicle followed by exposure with a proinflammatory cytokine cocktail (acute treatment (A), 1 h). Immunoblots (IB) shown are representative of 3 independent cell passages (A) and quantitation of the indicated protein levels (B-F). Tubulin was used as the loading control. Bars represent mean ± SEM. *P<0.05, **< 0.01, ****P<0.0001, ns= not significant. A vertical dashed line indicates the splicing of lanes from within the same gel exposure.

## 4. DISCUSSION

DOC2b deficiency and elevated CXCL10 is a feature of new-onset T1D islet β-cells [31], and elevated CXCL10 levels promote disease progression in the autoimmune mouse model [10–12]. Here, we report a novel role for DOC2b in attenuating cytokine stress-induced CXCL10 expression in human and mouse islets. We deduced that DOC2b attenuates cytokine-induced STAT-1 and IKKβ-IκBβ-NF-κB p65 signaling mechanisms — two known regulators of CXCL10 induction in cytokine-stressed β-cells [9, 44]. Marked attenuation of IKKβ and NF-κB p65 protein levels in DOC2b-human islets following cytokine challenge strengthen the concept that DOC2b is a suppressor of canonical NF-κB signaling and its downstream target CXCXL10 in stressed human islets. Moreover, we revealed a unique role for DOC2b in negatively regulating STAT-1 mRNA and protein abundances, and signaling in cytokine-challenged human islets, thereby establishing DOC2b’s dual protective action against CXCL10 in cytokine-stressed human islets. Emerging evidence suggest that crosstalk between ER stress and inflammation contributes to T1D pathogenesis [40, 41, 51]. In non-β cells, ER-stress induces NF-κB, independent of IKKα/β activation, but basal IKKβ, not IKKα level/activity, is required for the ER-stress-NF-κB crosstalk [52]. Here we revealed that DOC2b negatively regulates IKKβ (below basal level) and STAT-1 level in ER-stress sensitized-cytokine-stressed β-cells, indicating DOC2b’s possible mitigating action on ER-stress-NF-κB and STAT-1 crosstalk in β-cells. Taken together, these findings are consistent with a model wherein DOC2b attenuates IKKβ→IκBβ-NF-κB p65 and STAT-1 signaling and prevents cytokine-induced CXCL10 production in β-cells (Graphical Abstract).

Notably, we identified DOC2b as a key proximal modulator of IKKβ signaling. DOC2b is also an activator of STX4, and STX4 binds to/stabilizes IκBβ [20]. Therefore, DOC2b can directly and indirectly downregulate cytokine-induced NF-κB signaling and CXCL10 expression at two distinct nodes in this inflammatory signaling cascade. β-cell specific IKKβ mice were previously shown to spontaneously develop immune-mediated diabetes characterized by insulitis, hyperglycemia, and hypoinsulinemia [47], highlighting a critical role for IKKβ-NF-κB in immune-mediated diabetes. Intriguingly, successful remission of diabetes following transgene inactivation indicated modulation of IKKβ signaling as a potential therapeutic target for T1D [47]. Thus, DOC2b could potentially prevent IKKβ-mediated pathogenic signaling in β-cells.

Given that DOC2b activates STX4 in β-cells [25], we expected the anti-chemokine protective effects of DOC2b to use comparable mechanisms as STX4 in β-cells. Previously, STX4 was shown to dampen canonical NF-κB signaling via cytoplasmic retention of NF-κB inhibitory protein IκBβ, resulting in attenuation of CXCL9 expression and cytokine-induced apoptosis in β-cells [19, 20]. As with STX4, we observed increased levels of IκBβ, but not IκBα, in DOC2b-enriched cytokine-stressed human islets. In contrast to STX4, which is a downstream substrate for IKKβ [20], data here positions DOC2b as a putative upstream regulator of IKKβ in cytokine-stressed human islets and INS-1 832/13 β-cells. Indeed, our data showed that DOC2b and IKKβ can associate in INS-1 832/13 β-cells, consistent with the demonstration of association of recombinantly expressed and purified DOC2b and IKKβ proteins, a cell-free system. Thus, at least partially overlapping regulatory mechanisms for STX4 and DOC2b in modulating cytokine-stressed signaling in β-cells may exist.

Herein, DOC2b was also revealed to be a negative regulator of cytokine-induced STAT-1 expression in human islets. Indeed, a previous single-cell RNAseq study showed cytokine-stressed human islet β-cells have enhanced STAT-1 and CXCL10 [39]. Consistent with this, cytokine-responsive cis-regulatory elements are present in STAT-1, and NF-κB p65 in β-cells [53]. IL-1β and TNF-α signaling occurs via NF-κB [5, 6], and IFN-γ induces cytotoxicity via JAK-STAT [7, 8] regulated pathways - to our knowledge, DOC2b is the first suppressor common to both cytokine-induced STAT-1 and canonical NF-κB signaling pathway in β-cells. Previously in rat islets and rat β-cell line, IL-1β and IFN-γ were shown to cooperatively induce CXCL10 transcription via NF-κB p65 and STAT-1^Tyr701^; further, NF-κB p65 and STAT-1 are necessary for IFN-γ + IL-1β-induced CXCL10 expression [9, 44]. Here we showed that DOC2b could significantly block IL-1β-and IFN-γ-induced CXCL10 production in human islets, suggesting DOC2b dually regulated NF-κB and JAK-STAT pathways. Furthermore, we observed associations between DOC2b and IKKβ, NF-κB p65, and STAT-1 in β-cells that were acutely cytokine-stressed. We also bioinformatically predicted and biochemically confirmed DOC2b-IKKβ, and-STAT-1 protein-protein interactions, in cell-free systems. Indeed, we showed that DOC2b was a common modulator of STAT-1- and IKKβ-NF-κB p65-signaling in cytokine-stressed human islets and β-cells. Common repressors of STAT-1 and NF-κB signaling exist in other cell types [54, 55]. Currently, whether DOC2b prevents both STAT-1 and IKKβ signaling via modulation of their crosstalk or targets these two pathways independently remains unknown. Increased STAT- 1 gene expression in β-cell-specific IKKβ overexpressing mouse islets [47] suggests the existence of an inter-regulatory network of these two proteins, which could potentially be targeted by DOC2b. Intriguingly, DOC2b could prevent proinflammatory cytokine signaling upstream of JAK-STAT-1 and IKKβ by attenuating IL-1β and/or IFN-γ receptor signaling in β-cells. The precise DOC2b-regulated mechanisms linking STAT-1 and IKKβ downregulation in β-cells remains to be elucidated.

Which DOC2b domains/residues interact with IKKs, NF-κB p65, and STAT-1 and are required for CXCL10 regulation remain intriguing questions. In the rat INS-1 832/13 β-cell line, we showed that DOC2b’s C-terminal tandem C2AB domain was insufficient for mitigating cytokine-induced CXCL10 expression; in alignment, our bioinformatic prediction revealed DOC2b N-terminus involvement in DOC2b-IKK association. Our studies also showed Y301 phosphorylation as dispensable for mitigating cytokine-induced CXCL10 expression; bioinformatic predictions suggested against DOC2b Y301 involvement in DOC2b-STAT-1 association.

This study expands our knowledge of DOC2b in attenuating cytokine stress-induced CXCL10 expression in human islets, yet there are a few limitations. One limitation is in determining if this is specific to the human islet β-cells. Although islet β-cells are a known primary source of CXCL10 in T1D and NOD mice [56], islet α-cells, and immune cells are other sources of CXCL10 production in NOD mice and human T1D islets [56, 57]. It is possible that DOC2b was enriched in other human islet cell types, including α-cells, and infiltrating immune cells since DOC2b adenoviral expression was CMV-driven. We suggest that comparable results in both human islets and pure β-cell line point towards DOC2b’s protective effect against CXCL10 escalation, at least partially, in β-cells. Potential actions of DOC2b in cytokine-stressed α-cells and immune cells requires further elucidation. Secondly, human β-cells were shown to have a blunted cytokine-responsive gene expression versus mouse β-cells, which correlates with increased ribosomal protein [58]. Future single-cell analyses will be required to assess for any correlation between DOC2b protein levels, ribosomal protein gene expression, and cytokine-induced IKKβ/STAT-1. The third limitation is the absence of nuclear NF-κB p65 signal in our MLD-STZ treated mice models. As activation of classical NF-κB signaling is an early event of inflammation signaling [43, 59], we predict that the late-time point for pancreatic tissue harvest is the possible underlying reason. Additionally, future study is required to decipher if DOC2b’s direct binding with IKKβ, NF-κB p65 and STAT-1 is required for its anti-inflammatory function, and if yes, what is/are the responsible binding interfaces for these molecules in β-cells. Furthermore, DOC2b’s function in IKKβ/STAT-1 signaling in autoimmunity will need to be elucidated in vivo.

## 5. CONCLUSION

DOC2b enrichment protects against proinflammatory IKKβ→IκBβ/NF-κBp65- and STAT-1 signaling and attenuates CXCL10 induction. The DOC2b signaling hub may be a therapeutic target in β-cells to selectively block/reduce the immune storm in the β-cell milieu without the need for global immune suppression during T1D.

## Abbreviations

DOC2b: Double C-2 like domain beta
STX4: Syntaxin 4
MLD-STZ: multiple-low-dose-streptozotocin
NF-κB: Nuclear factor kappa-B
T1D: Type 1 diabetes
NOD mice: Nonobese diabetic mice
(h): Hours
(A): Acute
(C): Chronic
TBP: TATA Binding Protein
(GSIS): glucose-stimulated insulin secretion

## Acknowledgments

The authors would like to thank Dr. Peter Arvan (University of Michigan Medical Center) for helpful feedback during the review of this manuscript, Drs. Eunjin Oh and Rajakrishnan Veluthakal (City of Hope) for manuscript review, guidance in cell-free protein-protein interaction assay, and Erika McCown for technical support. The reported research includes work performed in the Multi-omics Mass Spectrometry & Biomarker Discovery Core, Light Microscopy/Digital Imaging Core, AR-DMRI Laboratory & Translational Service Centers, and the Integrative Genomics Core at the City of Hope. AR-DMRI Pancreatic Islet Cell & Tissue Processing Service Center (City of Hope) and the Integrated Islet Distribution Program supplied human islets for the study. The authors thank Dr. Chathurani S. Jayasena (City of Hope) for editing and providing helpful comments on the manuscript.

## CRediT authorship contribution statement

**Diti Chatterjee Bhowmick:** Writing – review & editing, Writing – original draft, Methodology, Investigation, Formal analysis, Data Curation, Resources, Funding acquisition, Conceptualization. **Miwon Ahn:** Writing – review & editing, Data Curation, Formal analysis. **Supriyo Bhattacharya:** Writing – review & editing, Writing –Methodology, Data Curation, Formal analysis. **Arianne Aslamy:** Writing – Review and editing, Resources. **Debbie C Thurmond:** Writing – review & editing, Writing – original draft, Supervision, Resources, Project administration, Funding acquisition, Conceptualization.

