## supplemental tables and figures for "DOC2b enrichment mitigates proinflammatory cytokine-induced CXCL10 expression by attenuating IKKβ and STAT-1 signaling in human islets"

**TABLE OF CONTENTS**

**Supplemental Information........................................................(ii-iii)**

**Supplemental Table 1……………………………………………………...(iv)**

**Supplemental Table 2……………………………………………….............(v)**

**Supplemental Table 3……………………………………………………...(vi)**

**Supplemental Table 4…………………………………………………….....(vii)**

**Supplemental Figure 1…………………………………………………….(viii)**

**Supplemental Figure 2…………………………………………………......(ix)**

**Supplemental Figure 3………………………………………………............(x)**

**Supplemental Figure 4……………………………………………………….(xi)**

**Supplemental Figure 5…………………………………………….........(xii)**

**Supplemental Figure 6…………………………………………………….(xiii)**

**Supplemental Figure 7…………………………………………………….(xiv)**

**Supplemental Figure 8……………………………………………………...(xv)**

**Supplemental Figure 9…………………………………………………….(xvi)**

**Supplemental Information.**

**A. Bioinformatics Analysis**: To investigate how DOC2b associates with IKKβ, NF-κB p65, and STAT-1, we used AlphaFold v3 to predict the corresponding protein complexes. AlphaFold (v3) predictions were performed using the online server at https://alphafoldserver.com. For each complex, the top-ranked structure given by Alphafold was used for analysis. DOC2b and other protein sequences were obtained from UniProt. In each case, the isoform designated as canonical in Uniprot was selected. The following Uniprot accession IDs were used: DOC2B: Q14184, munc18-1: P61764, munc18c: O00186, IKKα: O15111, IKKβ: O14920, NF-κB p65: Q04206, STAT1: P42224. Full-length DOC2b sequence was used in all instances except the DOC2b double C2 domains were used for determining munc18 association predictions. Our AlphaFold analysis method was verified by capturing the known association of DOC2b with munc18-1 and phosphorylated munc18c (Suppl Fig. 5A-B). Munc18-1 and munc18-c are known to associate with the double C2 domain of DOC2b [1]. Previously, we showed that in β-cells, DOC2b, munc18c, and munc18-1 form a heterotrimeric complex where DOC2b acts as a scaffold bringing the two munc18 proteins together, facilitating GSIS [2]. Based on AlphaFold, the two munc18 proteins were predicted to associate to opposite faces of the C2 domains of DOC2b with minimal overlap with each other, thereby supporting our previous observation. Also, Phospho-Y219 on munc18c, the residue known to be critical for association with DOC2b [3], was predicted to be at the protein-protein interface and engaged in a crucial salt-bridge with K321 of DOC2b. This explains our previous observation about the importance of this phospho-site in recognizing DOC2b.

We next proceeded to predict the association of DOC2b with the newly found partners in this work, namely IKKα/β, NF-κB p65 and STAT-1. For IKKα/β, the DOC2b double C2 domain was consistently predicted to associate near the IKKα/β kinase domains (Suppl Fig. 5C-E); moreover, several serine residues necessary for IKKα/β complex activation were located near the DOC2b association site [4]. Therefore, DOC2b may prevent IKKα/β activation via steric hindrance. A short N-terminal α-helical segment of DOC2b was inserted between the two IKKα/β monomers (Suppl Fig. 5C-E), potentially interfering with IKK activation by NEMO [4]. Also predicted DOC2b bound IKKα dimer structure showed a closer confirmation vs IKKα dimer crystal structure [5], indicating an inactive conformation of IKKα while bound to DOC2b (Suppl Fig. 5F). Thus, double C2 and N-terminal domains of DOC2b likely inhibit IKKα/β. NF-κB p65 and STAT-1 associate substantially with the DOC2b double C2 domains. While the main association interface with NF-κB p65 involved the C2A domain, we found a tight association between the C2B domain and a short C-terminal α-helical segment of NF-κB p65 (Suppl Fig. 5G). The DOC2b C2B domain is associated with the known STAT1 dimer interface, suggesting that DOC2b could inhibit STAT1 dimerization and activation (Suppl Fig. 5H) [6]. However, all the above-discussed bioinformatics predictions need to be verified in future biochemical studies. Taken together, these observations support and rationalize the structural basis of our experimental evidence regarding the association of DOC2b with these proteins.

**B. Glucose stimulated insulin secretion (GSIS):** Glucose stimulated insulin secretion from vector and DOC2b-GFP transduced INS-1 832/13 β-cells, with /without chronic cytokine (C) exposure was measured under conditions of static incubation using Krebs-Ringer bicarbonate HEPES buffer (KRBH) [7]. Briefly, INS-1 832/13 β-cells were equilibrated in low-glucose (2.5 mmol/L) and low-serum (2.5%) culture medium overnight. The KRBH was oxygenated with $O_2$ gas, after which 0.1% w/v fatty acid–free BSA was added (Sigma-Aldrich). Before insulin secretion assays, cells were incubated in KRBH buffer containing 1% radioimmunoassay-grade BSA for 1 h, followed by stimulation with 20 mmol/L glucose (30 min). 2.5 mmol/L glucose was used for basal conditions. A Rat High Range Insulin ELISA kit (Alpco, Salem, NH) was used to quantify insulin.

**References.**

**Supplemental Table 1. Human islet donor information**

| RRID | Sex | Age (year) | BMI | HbA1C | T2D | Used For |
|---|---|---|---|---|---|---|
| SAMN12274306 | M | 37 | 25.30 | 5.20 | No | Fig. 2A(i-iii); Suppl. Fig. 1A |
| SAMN12500521 | M | 52 | 29 | 5.10 | No | Fig. 2A(i-iii); Suppl. Fig. 1A |
| SAMN12598151 | M | 51 | 31.50 | 5.20 | No | Fig. 2A (i-iii); Suppl. Fig. 1A |
| SAMN12129273 | F | 30 | 35.70 | 4.50 | No | Fig. 2A (i-iii); Fig. 3A; Suppl. Fig. 1A; Suppl. Fig. 4A |
| SAMN12292085 | M | 20 | 35.80 | 4.60 | No | Fig. 2A (i-iii); Fig. 3A; Suppl. Fig. 1A; Suppl. Fig. 4A |
| SAMN12339206 | M | 45 | 32.90 | 5.40 | No | Fig. 2A (i-iii); Fig. 3A; Suppl. Fig. 1A; Suppl. Fig. 4A |
| SAMN31242270 | M | 52 | 37.50 | 5.30 | No | Fig. 3A; Suppl. Fig. 1A; Suppl. Fig. 4A |
| SAMN30686018 | M | 39 | 32.40 | 4.30 | No | Fig. 2A (iv-v) |
| SAMN28867622 | M | 36 | 29.60 | 5.40 | No | Fig. 2A (iv-v) |
| SAMN14132340 | M | 31 | 27 | 5.20 | No | Fig. 2A (iv-v) |
| SAMN37973608 | F | 55 | 31.70 | 5.90 | No | Fig. 2A (iv-v) |
| SAMN37159756 | M | 34 | 33.3 | 5.20 | No | Fig. 6 A-B; Fig. 7A; Suppl. Fig. 7, 8A |
| SAMN38428316 | M | 51 | 25.60 | 5.90 | No | Fig. 6 A-B; Fig. 7A; Suppl. Fig. 7, 8A |
| SAMN38518088 | F | 57 | 30.10 | 5.80 | No | Fig. 6A-B; Fig. 7A; Suppl. Fig. 7, 8A |
| SAMN37350251 | F | 43 | 29.89 | 5.20 | No | Fig. 6C-D |
| SAMN37638596 | M | 34 | 27 | 5.5 | No | Fig. 6C-D |
| SAMN38117428 | M | 47 | 28.30 | 5.30 | No | Fig. 6C-D |
| SAMN38750927 | M | 34 | 24.40 | 5.40 | No | Fig. 4B; Fig. 7B-C; Suppl. Fig. 2A, B |
| SAMN36705973 | F | 36 | 30.10 | 4.80 | No | Fig. 4B; Fig 7B-C; Suppl. Fig. 2A, B |
| SAMN40725442 | M | 44 | 24.4 | 5.60 | No | Fig. 4B; Fig 7B-C; Suppl. Fig. 2A, B |
| SAMN42008301 | M | 36 | 30 | 5 | No | Fig. 4B |
| SAMN41218531 | M | 38 | 29 | 5.20 | No | Fig. 4B |

### Supplemental Table 2. qPCR primer list

| Species | Gene | Forward primer (5' to 3') | Reverse primer (5' to 3') |
|---|---|---|---|
| Human | CXCL10 | AGCAAGGAAAGGTCTAAAAGATCTCC | GGCTTGACATATACTCCATGTAGGG |
| | CXCL9 | ACTATCCACCTACAATCCTTGAAAGAC | TCACATCTGAATCTGGGTTTAG |
| | DOC2b | TGGTGTGGTTCTGGGCATCCACG | TGGGAGCTCGCTGGTGAGCGTG |
| | ATF4 | Qiagen: QT00366233 | Qiagen: QT00366233 |
| | STAT-1 | 5'-ATGGCAGTCTGGCGGCTGAATT-3' | 5'-CCAAACCAGGCTGGCACAATTG-3 |
| | GAPDH | GTCTCCTCTGACTTCAACAGCG | ACCACCCTGTTGCTGTAGCCAA |
| Rat | CXCL10 | GCAAGTCTATCCTGTCCGCAT | GGGTAAAGGGAGGTGGAGAGA |
| | CXCL9 | CAAGGCACATTCCACTACAA | CCTTGCTGAATCTGGGTCTA |
| | DOC2b | CCAGCAAGGCAAATAAGCTC | GTTGGGTTTCAGCTTCTTCA |
| | CHOP | Qiagen: QT01080772 | Qiagen: QT01080772 |
| | GAPDH | Qiagen: PPR06557B | Qiagen: PPR06557B |
| Mouse | CXCL10 | Qiagen: PPM02978E | Qiagen: PPM02978E |
| | DOC2b | CAGTCTGCTCTATGACCAGGAG | ATCAGCCAGTCCATTGTGGTCC |
| | GAPDH | Qiagen: QT10658692 | Qiagen: QT10658692 |

Fifty nanograms of RNA were used for quantitative real-time PCR using the one-step qPCR kit (Qiagen) with SYBR. qPCR conditions: cDNA synthesis at 50°C for 30 min and 95°C for 2 min hold, then 40 cycles of 95°C for 15 sec, 58°C for 30 sec and 72°C for 15 sec.

### Supplemental Table 3. Primary antibody list

| # | Antibody Name | Vendor | Cat# | Dilution used | Purpose |
|---|---|---|---|---|---|
| 1. | Rabbit anti-DOC2b | In-house | | 1:1000 | IB |
| 2. | DDK (FLAG) mouse mAb | Origene Technologies | TA180144 | 1:2000 | IB |
| 3. | Rabbit anti-GFP | Abcam | Ab290 | 1:2000 | IB |
| 4. | Living colors, Mouse anti-GFP | Takara Bio | 632381 | 1:5000 | IB |
| 5. | Mouse anti-Tubulin | Millipore Sigma | T5168 | 1:30,000 | IB |
| 6. | Guinea pig anti-Insulin | Thermo Fisher | PA126938 | 1:50 | IHC |
| 7. | Rabbit anti-CXCL10 | Invitrogen | PA5-115067 | 1:1000 (IB)<br>1:100 (IHC) | IB, IHC |
| 8. | Mouse anti-p-Stat1 (A-2) (Tyr 701) | Santa Cruz Biotech | sc-8394 | 1:500 | IB |
| 9. | Mouse anti-Total-STAT-1 | Santa Cruz Biotech | Sc-464 | 1:500 | IB |
| 10. | Total-STAT-1 Rabbit mAb | Cell Signaling | 14994S | 1:1000 | IB (INS-1 832/13 β-cell) |
| 11. | p-IKKα/β (ser 176/180) Rabbit mAb | Cell Signaling | 2697S | 1:1000 | IB |
| 12. | Total-IKKβ Rabbit mAb | Cell Signaling | 8943S | 1:1000 | IB |
| 13. | Total-IKKα Rabbit mAb | Cell Signaling | 61294 | 1:1000 | IB |
| 14. | IκBβ (7B4) Mouse mAb | Cell Signaling | 8635S | 1:1000 | IB |
| 15. | IκBα Rabbit mAb | Cell Signaling | 4814S | 1:1000 | IB |
| 16. | NF-κB p65 Rabbit mAb | Cell Signaling | 8242S | 1:1000 (IB), 1:100 (IHC) | IB, IHC |
| 17. | TBP Antibody | Cell Signaling | 8515S | 1:1000 | IB |
| 18. | Rabbit Cleaved Caspase 3 | Cell Signaling | 9661S | 1:1000 | IB |
| 19. | Rabbit Phospho-eIF2α | Cell Signaling | 9721S | 1:1000 | IB |
| 20. | Total eIF2α mAb | Cell Signaling | 2103S | 1:1000 | IB |
| 21. | Rabbit anti-Rho-GDI | Santa Cruz Biotech | Sc-360 | 1:5000 | IB |
| 22. | ChromoTek DYKDDDDK Fab-Trap® Agarose | Proteintech | ffa | 25μl/reaction | Co-IP |
| 23. | ChromoTek GFP-Trap® Magnetic Agarose | Proteintech | gtma | 25μl/reaction | Co-IP |

IB: Immunoblot/western blot; IHC: Immunohistochemistry; co-IP: co-Immunoprecipitation.

**Supplemental Table 4. Potential DOC2B binding proteins identified by mass spectrometry**

| | | | *Exclusive Unique Peptide Count* | |
|---|---|---|---|---|
| # | **Protein name** | **Alternate ID** | **DOC2b-GFP Basal** | **GFP Basal** |
| * | Double C2-like domain-containing protein beta | DOC2b | 37 | 0 |
| 1 | Nuclear factor NF-kappa-B p105 subunit | NF-κB1 | 6 | 0 |
| 2 | Nuclear factor NF-kappa-B p100 | NF-κB2 | 18 | 0 |
| 3 | Inhibitor of nuclear factor kappa-B kinase subunit beta | IKKβ | 9 | 0 |
| 4 | Inhibitor of nuclear factor kappa-B kinase subunit alpha | IKKα | 10 | 0 |
| 5 | Signal transducer and activator of transcription 1 | STAT-1 | 14 | 0 |
| 6 | Signal transducer and activator of transcription 3 | STAT-3 | 12 | 0 |
| 7 | Signal transducer and activator of transcription 5A | STAT-5A | 7 | 0 |

*DOC2B-GFP was successfully expressed and captured (positive control). The listing of DOC2b binding proteins captured by mass spectrometry includes only those with >3 peptides.

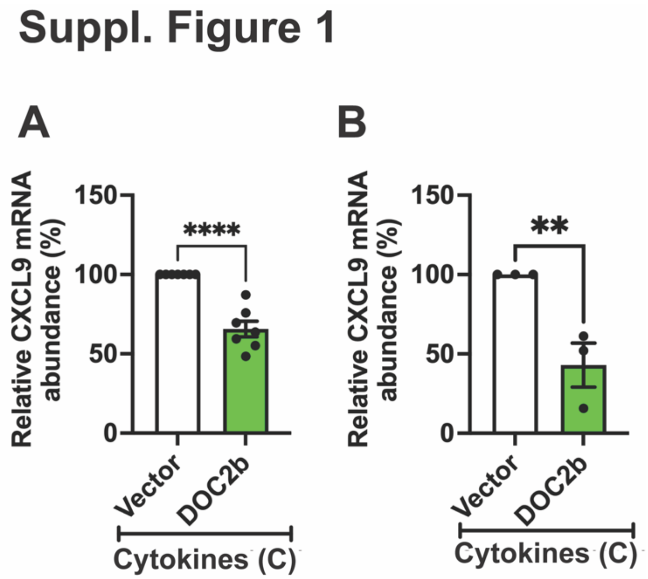


**Suppl. Figure 1. DOC2b overexpression protects against proinflammatory cytokine-induced CXCL9 gene expression in human islets and INS-1 832/13 β-cells.** (A) DOC2b- or control (vector)-transduced non-diabetic human islets were exposed to a proinflammatory cytokine cocktail (chronic treatment (C), ≥ 8h) and qPCR quantitation of CXCL9; n=7 independent human donors/group. (B) DOC2b-GFP- or vector control (GFP)-transduced INS-1 832/13 β-cells were exposed to a proinflammatory cytokine cocktail (chronic treatment (C), ≥ 8h) and qPCR quantitation of CXCL9; n =3 independent cell passages. Bars represent the mean ± SEM. **P< 0.01, ****P<0.0001.

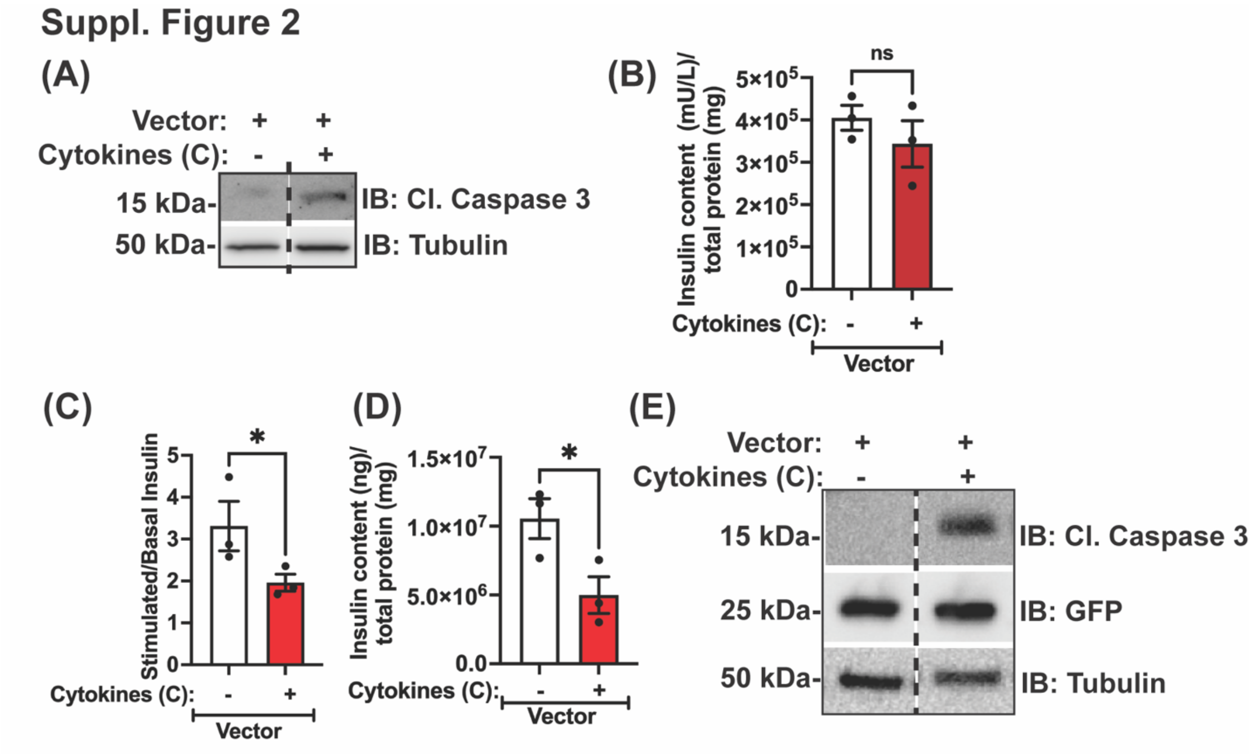


**Suppl. Figure 2. Validation of chronic cytokine treatment in human islets and INS-1 832/13 β-cells.** (A-B) Vector-transduced non-diabetic human islets were exposed to a proinflammatory cytokine cocktail (C) for ≥ 8h. (A) Immunoblot analysis (IB) representative of 3 independent human donor islet batches for the stated proteins. Loading control: tubulin. (B) Quantification of the insulin content. (C-E) Vector-transduced INS-1 832/13 β-cells were exposed to a proinflammatory cytokine cocktail (C) for ≥ 8h. (C) Glucose stimulated insulin secretion (GSIS) assay using 2.5 mM (Basal) and 20 mM (stimulated, 30 minutes) glucose. (D) Quantification of the intracellular insulin content. (E) Immunoblot analysis (IB) is representative of 3 independent cell passages for the designated proteins. Loading control: tubulin. Bars represent the mean ± SEM. *P< 0.05, ns= not significant. A vertical dashed line indicates the splicing of lanes from within the same gel exposure.

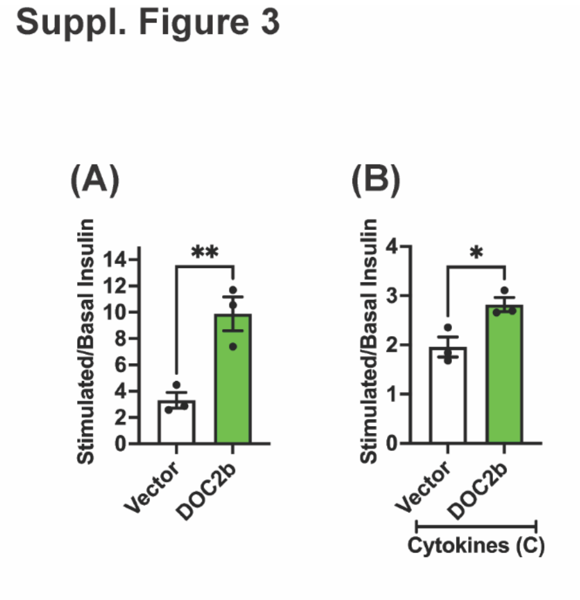


**Suppl. Figure 3. DOC2b overexpression boosts GSIS in cytokine-challenged INS-1 832/13 β-cells.** (A-B) Vector (GFP) and DOC2b-GFP-transduced INS-1 832/13 β-cells were incubated with or without proinflammatory cytokine cocktail (C) for ≥ 8h. Glucose stimulated insulin secretion (GSIS) assay using 2.5 mM (Basal) and 20 mM (stimulated, 30 minutes) glucose. Insulin released was quantified using ELISA. Bars represent the mean ± SEM. *P< 0.05, **P< 0.01.

### Suppl. Figure 4

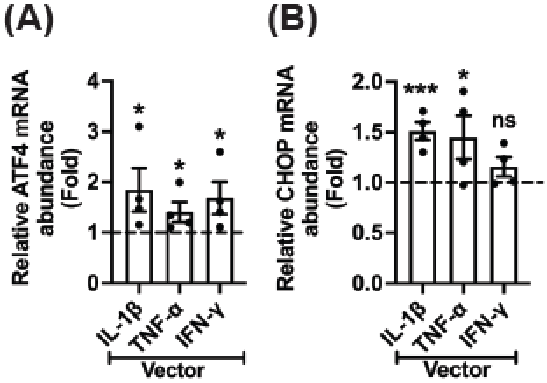


**Suppl. Figure 4. Validation of ER-stress induction in human islets and INS-1 832/13 β-cells challenged with individual cytokines.** Control (vector)-transduced primary non-diabetic human islets (A, n= 3 donors) or INS-1 832/13 β-cells (B, n= 3 passages) were chronically exposed to a species-specific cytokines IL-1β/TNF-α/IFN-γ for >8h. Quantitation of ER-stress marker ATF4 (A) and CHOP (B). Bars represent the mean ± SEM. *P<0.05, ***P<0.001, ns= non-significant compared to no cytokine control. Horizontal dashed lines represent fold gene expression without cytokine exposure.

### Suppl. Figure 5

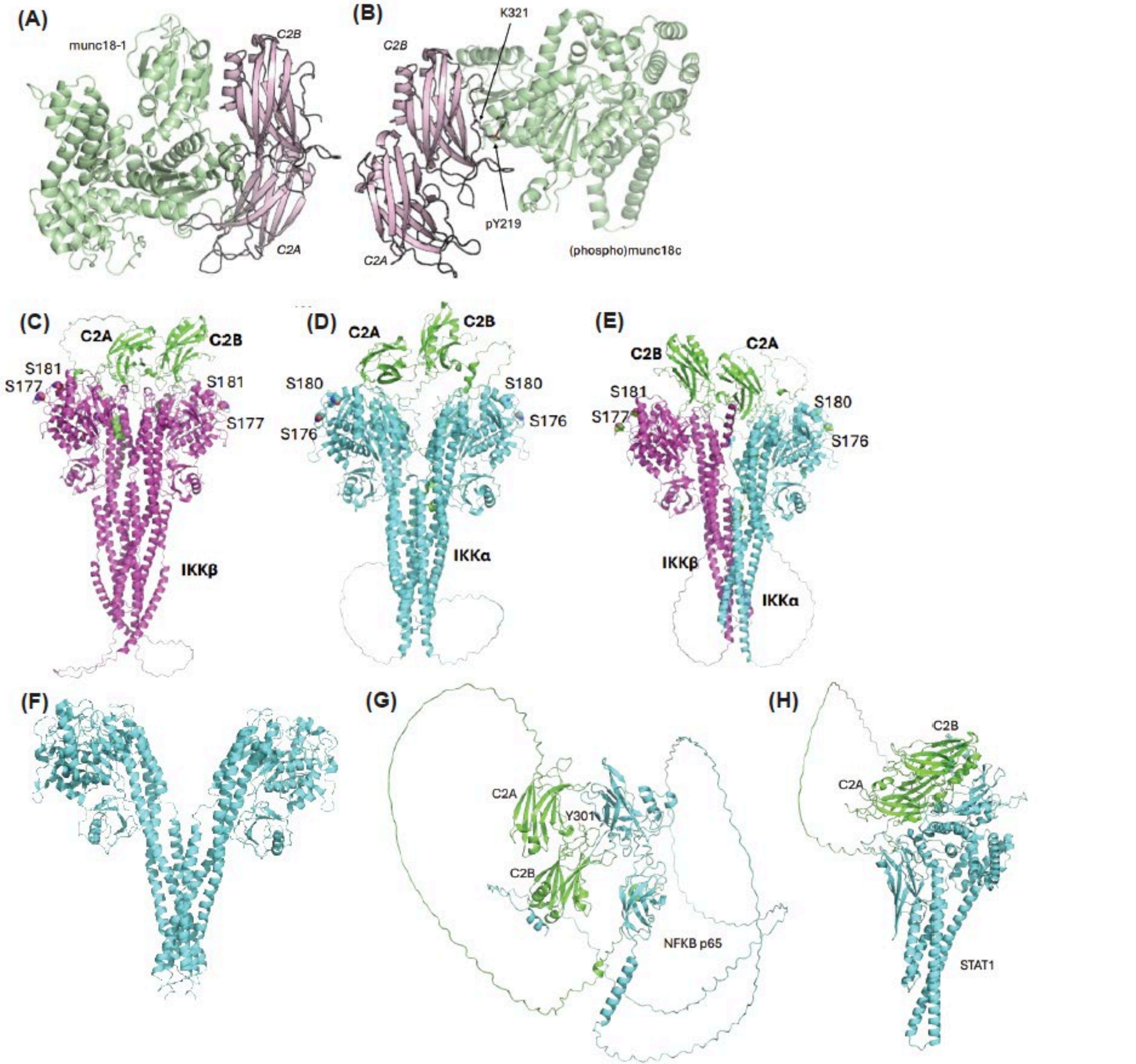


**Suppl. Figure 5. Predicted DOC2b association with Munc18, IKKα, IKKβ, NF-κB p65 and STAT-1 using AlphaFold.** (A-B) Predicted complexes of DOC2B C2 domains with (A) Munc18-1, and (B) (phospho)-Munc18c. Pink: DOC2B. Green: Munc18 molecules. (C-E) Predicted DOC2B complexes with IKKα homodimer (C), IKKβ homodimer (D), and IKKα/ IKKβ heterodimer (E). Green: DOC2B. Cyan: IKKα. Magenta: IKKβ. The active site serine residues whose phosphorylation is critical for activating the IKK proteins are shown as spheres and labeled. (F) Crystal structure of human IKKα dimer (PDB id 5EBZ). (G-H) Predicted complexes of DOC2B with NF-κB p65 (G) and STAT-1 (H). Green: DOC2B.

### Suppl. Figure 6

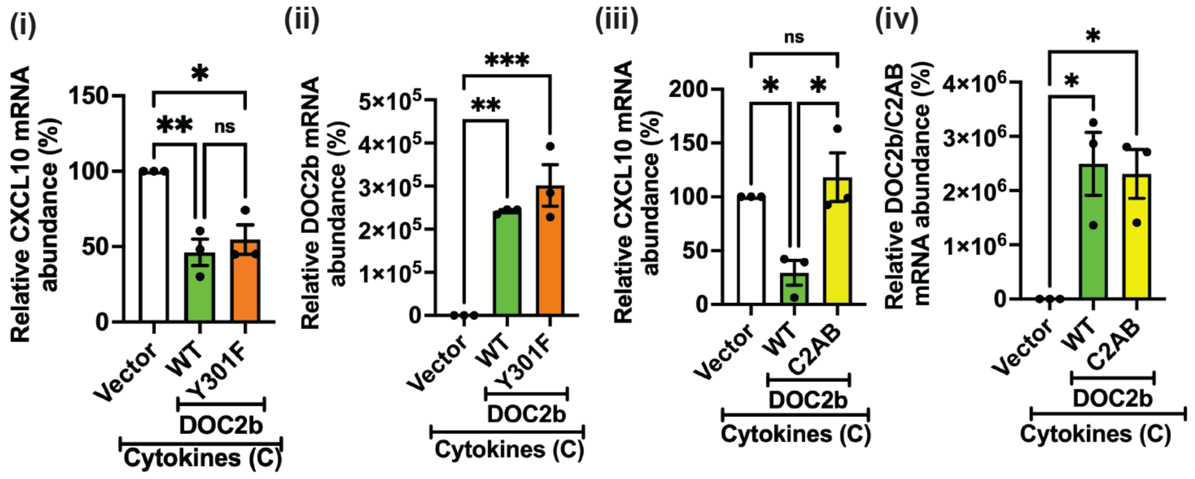


**Suppl. Figure 6. DOC2b Y301 phosphorylation is not required, and C2AB peptide is not sufficient, to reduce cytokine-induced CXCL10 gene expression in INS-1 832/13 β-cells.** DOC2b-GFP$^{WT/Y301F}$ (i-ii) or DOC2b-MYCDDK$^{WT/C2AB}$ (iii-iv) or vector-control-transduced INS-1 832/13 β-cells were exposed to a cytokine cocktail (chronic treatment (C), ≥ 8h). (i, iii) qPCR quantitation of CXCL10 (ii-iv), and DOC2b$^{WT/Y301F/C2AB}$ (iii); n=3 independent cell passages. Bars represent the mean ± SEM *P<0.05, **P< 0.01, ***P<0.001, ns= not significant.

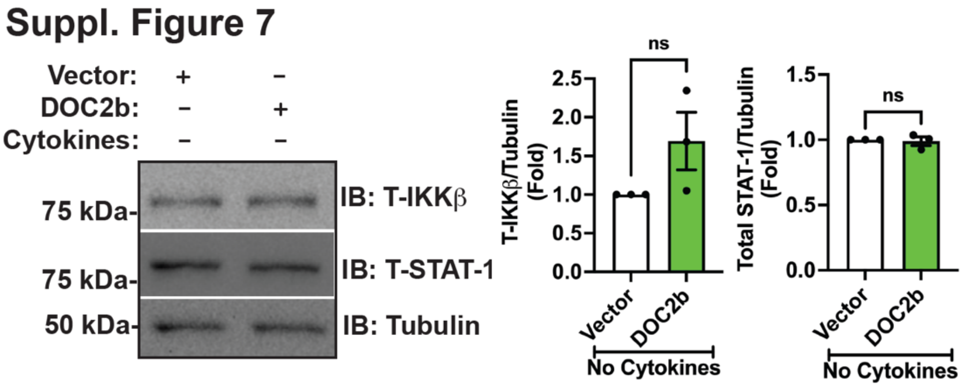


**Suppl. Figure 7. DOC2b overexpression did not reduce total IKKβ and STAT-1 levels without cytokine treatment in human islets.** Non-diabetic human islets were transduced with DOC2b- or control (vector)-adenoviruses. Immunoblot analysis (IB) representative of 3 independent human donor islet batches for the designated proteins, and bar graph quantitation of the indicated protein levels. Tubulin was used as the loading control. Bars represent the mean ± SEM (n=3 independent donors). ns= not significant.

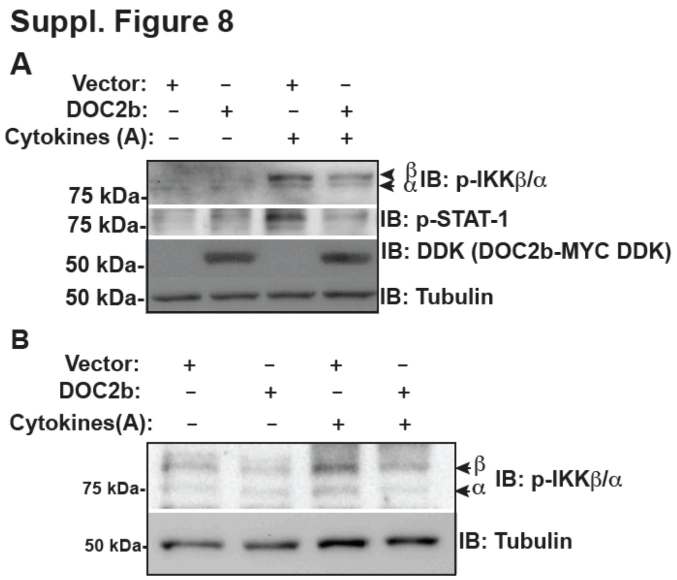


**Suppl. Figure 8. Validation of action of cytokines on activation of IKK complex and STAT-1 in human islets and INS-1 832/13 β cells.** (A) DOC2b- or control (vector)-transduced non-diabetic human islets were treated with a proinflammatory cytokine cocktail (acute treatment (A), 1h) and Immunoblot analysis (IB) for the designated proteins; representative of n=3 independent human donor islet batches. Tubulin was used as the loading control. (B) DOC2b-GFP- or vector control (GFP)- transduced INS-1 832/13 β-cells were treated with a proinflammatory cytokine cocktail (acute treatment (A), 1 h) and Immunoblot analysis (IB) for the designated protein; representative of n= 3 independent cell passages. Tubulin was used as the loading control.

Suppl. Figure 9

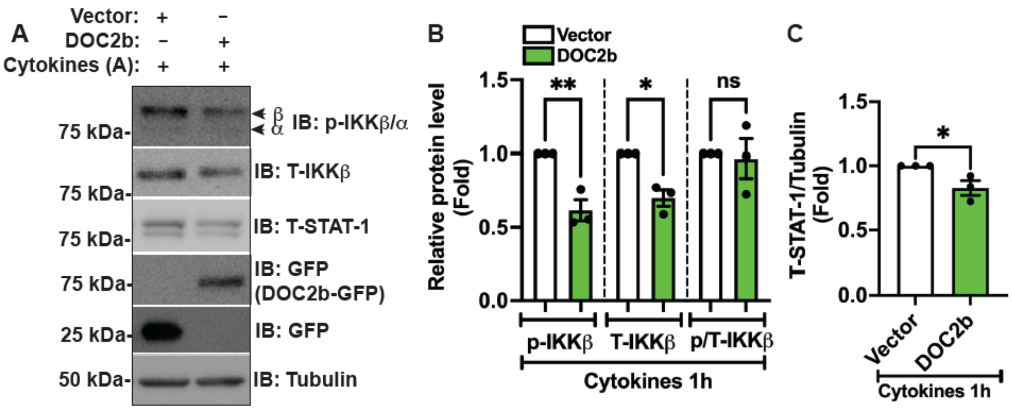


**Suppl. Figure 9. DOC2b enrichment reduces IKKβ and STAT-1 protein levels in INS-1 832/13 β-cells following acute cytokine treatment.** DOC2b-GFP- or vector control (GFP)-transduced INS-1 832/13 β-cells were treated with a proinflammatory cytokine cocktail (acute treatment (A), 1 h). Immunoblots (IB) shown are representative of 3 independent cell passages (A) and quantitation of the indicated protein levels (B-C). Tubulin was used as the loading control. Bars represent mean ± SEM. *P<0.05, **< 0.01, ns= not significant.
